# Polysome-CAGE of TCL1-driven chronic lymphocytic leukemia revealed multiple N-terminally altered epigenetic regulators and a translation stress signature

**DOI:** 10.1101/2022.02.15.480558

**Authors:** Ariel Ogran, Tal Havkin-Solomon, Shirley Becker-Herman, Keren David, Idit Shachar, Rivka Dikstein

## Abstract

The transformation of normal to malignant cells is accompanied by substantial changes in gene expression programs through diverse mechanisms. Here we examined the changes in the landscape of transcription start sites (TSSs) and alternative promoter (AP) usage and their impact on the translatome in TCL1-driven chronic lymphocytic leukemia (CLL). Our findings revealed a marked elevation of APs in CLL cells from Eµ-Tcl1 transgenic mice, which are particularly enriched with intragenic promoters that generate N-terminally truncated or modified proteins. Intragenic promoter activation is mediated by (i) loss of function of ‘closed chromatin’ epigenetic regulators due to the generation of inactive N-terminally modified isoforms or reduced expression; (ii) upregulation of transcription factors, including c-Myc, targeting the intragenic promoters and associated enhancers. Exogenous expression of Tcl1 in MEFs is sufficient to induce intragenic promoters of epigenetic regulators and promote c-Myc expression. We further found a dramatic translation downregulation of transcripts bearing CNY cap-proximal tri-nucleotides, reminiscent of cells undergoing metabolic stress. These findings uncovered the role of Tcl1 oncogenic function in altering promoter usage and mRNA translation in leukemogenesis.

## Introduction

Chronic Lymphocytic Leukemia (CLL) CLL accounts for 25% to 30% of all leukemias in the Western countries, with incidence rates ranging from 3.65 to 6.75 cases per 100,000 population per year (1, 2). CLL is characterized by an outgrowth of malignant CD5 positive B cells, mainly residing in the peripheral blood, bone marrow, and lymphoid organs, and by a high biological heterogeneity reflected in clinically different outcomes including disease progression, therapy response, and relapse (3, 4). The hallmark of the CLL disease is mainly decreased apoptosis resulting from overexpression of anti-apoptotic proteins and resistance to apoptosis in vivo. The TCL1 oncogene is highly expressed in CLL cells, and its expression level correlates with the CLL aggressiveness (5). TCL1 was initially discovered as a gene involved in the rearrangement of the T cell leukemia/lymphoma 1 locus at 14q32.1, the most common chromosomal aberrations detected in mature T-cell leukemias (6). The central role of TCL1 in CLL was demonstrated in transgenic mice overexpressing the human TCL1 gene under the control of the immunoglobulin promoter and enhancer (Eµ-Tcl1) (7). These mice develop normally into adulthood but develop CLL that is highly similar to the human disease and is characterized by CD5+ cells, which accumulate in the spleens, livers, and lymph nodes. The association of high TCL1 expression and malignant transformation is linked to its involvement in several major oncogenic pathways, including the enhancement of AKT activity(8–11); interaction with the DNA repair ataxia-telangiectasia-mutated (ATM) factor (12), and the transcription factor CREB (13) as well as in contributing to accelerated tumorigenic and pro-survival NF-κB signaling (12). Inhibition of activator protein 1 (AP-1) transcriptional activity via interaction of TCL1 with the AP-1 complex represents another mechanism to antagonize the expression of pro-apoptotic factors (12, 14). TCL1 was also shown to contribute to epigenetic reprogramming via interacting with the de novo DNA methyltransferase 3A (DNMT3A) and reducing its enzymatic activity (15). Considering the central role of TCL1 in tumor initiation, progression, and maintenance, it is reasonable that its oncogenic functions involve additional, yet to be discovered, oncogenic activities.

The transcription initiation of a specific gene can occur from more than one position in the DNA, creating several mRNA isoforms. Alterations in transcription start site (TSS) selection and alternative promoter (AP) usage increase transcriptome diversity and regulation. For example, the level of transcription initiation can vary between different TSSs under different growth conditions, in response to a specific signal, or different cell types and tissues. In addition, mRNA isoforms with different 5’ leaders can vary in their translation efficiency or half-lives. Likewise, AP usage can lead to the generation of protein isoforms that differ in their N-termini and as a result of different or even opposite biological functions. Recent large-scale promoter analysis in hundreds of human and mouse primary cell types shed light on the prevalence of AP usage in mammals (16) as well as in multiple cancer types (17). Several studies have examined the translation and stability of transcript isoforms of the same gene and the contribution of AP usage to the translational response to stress (18–21). However, presently little is known about the impact of TSSs and AP usage changes on the protein content of cells undergoing cancer progression. Considering that even transient cellular stress is associated with dramatic changes in the landscape of TSSs and AP usage that are coordinated with translation (21), it is reasonable to assume that such alterations can affect translatability in cells undergoing oncogenic transformation. Some of the new cancer-specific protein isoforms generated by AP usage can help maintain the transformed phenotype. Presently, the vast majority of studies measured global changes in mRNA abundance and transcript translatability in cancer cells, while the potential effect of variations in TSSs selection and AP usage on translation was hardly addressed.

This study examined the changes in TSS selection and AP usage and their impact on the translatome during malignant transformation of B cells by the TCL1 oncogene. Using healthy and Eµ-Tcl1 transgenic mice, we compared the TSSs landscape of normal and CLL B cells. Our findings revealed a marked elevation of APs in malignant CLL B cells with a particularly high prevalence of intragenic promoters that are predicted to generate N-terminally truncated or N-terminally modified proteins. The induced cryptic promoters are driven by nearby enhancers and up-regulation of their cognate transcription factors, including the c-Myc oncogene. A promoter shift in the opposite direction accounts for the induced expression of the immune checkpoint ligand PD-L2. Notably most of the intragenic-generated transcripts are also efficiently translated. These remarkable changes of promoters and TSSs are linked to the loss of function of many ’closed chromatin’ epigenetic regulators via induction of inactive isoforms by promoter shifts or by reduced expression, resulting in a feed-forward loop. The activation of intragenic cryptic promoters and the c-Myc oncogene elevation are mediated, at least in part, by the Tcl1 oncogene itself and are therefore intrinsic to the CLL B cells. We further found a dramatic and specific translation downregulation of transcripts bearing CNY cap-proximal tri-nucleotides, reminiscent of cells undergoing energy stress. These findings explored the contribution of 5’ end mRNA isoforms and their associated translational changes during cellular transformation and uncovered novel Tcl1 oncogenic pathways.

## Results

### Global analysis of TSSs revealed extensive induction of promoter shifts in CLL B cells

To address the potential contribution of AP to CLL we used the Eµ-Tcl1 transgenic mice overexpressing the human TCL1 gene under the control of the immunoglobulin heavy chain (IgH) variable region promoter (VH) and immunoglobulin heavy chain enhancer (Eµ-Tcl1) (7). Like the human CLL, Eµ-Tcl1 mice accumulate transformed CD5+ B cells in the spleens, livers, and lymph nodes during adulthood, reproducing CLL (13). Splenocytes from healthy and Eµ-Tcl1 mice were subjected to affinity-based capture using CD19 magnetic beads to isolate B cells and CLL B cells, respectively. Total RNA was extracted and subjected to Cap Analysis of Gene Expression (CAGE) (22, 23) library preparation and Illumina deep sequencing of the first 27 nucleotides CAGE tags from the 5’ ends (Fig. 1A). An average of 10.625 million CAGE tags per sample were aligned to a total of 2.67 million CAGE-derived TSSs (CTSSs) that were mapped and clustered to a consensus set of 28,616 promoters that correspond to 11,515 genes. Using this high-throughput data from two independent biological replicates, we determined the global landscape of TSSs and their relative abundance in healthy and CLL B cells. Metagene analysis of both WT and Eµ-Tcl1 CAGE data showed almost perfect alignment to the annotated TSS atlas of the FANTOM5 project (24) (Fig. 1B), indicating the proper calculation of the TSS positions at a single-base resolution. As expected, in both WT and Eµ-Tcl1 samples, most of the TSSs with the highest expression level were mapped to the core promoter region (Fig. 1C). A significant fraction of the TSSs was found within introns and intergenic regions. Since the CAGE method captures all capped RNAs, this data also detect enhancer RNAs (eRNAs), a class of short non-coding RNAs resulting from a balanced bidirectional transcription from enhancers that are unstable and non-polyadenylated (25, 26) (examples of an enhancer and super-enhancer are shown in Figure 1 – figure supplement 1A and B). We examined the possibility that part of the intronic and intragenic CAGE reads are derived from active eRNAs. Based on the criteria of balanced bidirectional read mapping, we calculated the frequencies of eRNAs in introns and intergenic regions and found that most of the intronic TSSs (85% in Eµ-Tcl1 and 79% in WT) and a significant fraction of the intragenic TSSs (62% in Eµ-Tcl1 and 50% in WT) are indeed derived from eRNAs (Fig. 1C). The ability of CAGE to detect lowly expressed genes makes it crucial to reduce the mean-variance relationship of count data prior to differential expression (DE) analysis. A “blind” analysis of the Variance Stabilizing Transformation (VST) was performed and summarized by a Principle Component Analysis (PCA). PCA1 explained 85% of the variance and distinguished between WT and Eµ-Tcl1, where WT samples are tightly clustered and Eµ-Tcl1 are separated by 13% of the variance, which is explained by PCA2 (Figure 1 – figure supplement 1C). As no systemic effect on samples was found, we decided to fit a DE model without using a correction factor. Out of the 28,616 consensus TSSs, we found in Eµ-Tcl1 3,337 upregulated and 1,829 downregulated TSSs (Fig. 1D), suggesting an overall upregulation of many promoters in CLL. To test the contribution of eRNA to the differentially expressed TSSs in Eµ-Tcl1, we used an established method to predict eRNA-TSS interactions based on genomic distance and expression-based correlation(27, 28). We found 255 upregulated eRNAs, of them 106 were positively correlated to 203 upregulated TSSs, while out of 179 downregulated eRNAs, 86 were positively correlated to 331 downregulated TSSs (Fig. 1D). Lengths of the TSS clusters were calculated by the interquartile range (IQR) of highly expressed (> 10 TPM) TSSs. TSS clusters lengths holding 10%-90% of pooled TSSs expression were found to distribute into sharp and broad promoters bimodally (Figure 1 – figure supplement 1D). Consistent with known promoter architecture (22, 29), general TSSs are characterized by YR at the (-1) and (+1) positions (Figure 1 – figure supplement 1E). Promoters with broad TSS distributions tend to be surrounded by CpG islands, while promoters with sharp TSSs are defined by the presence of a TATA box around position -30 (Figure 1 – figure supplement 1E). Separately plotted LOGOs of differentially expressed TSSs, showed enrichment of poly-pyrimidine sequence in the upregulated sharp TSSs at the positions of +2 to +12 (Fig. 1E).

**Figure 1:**
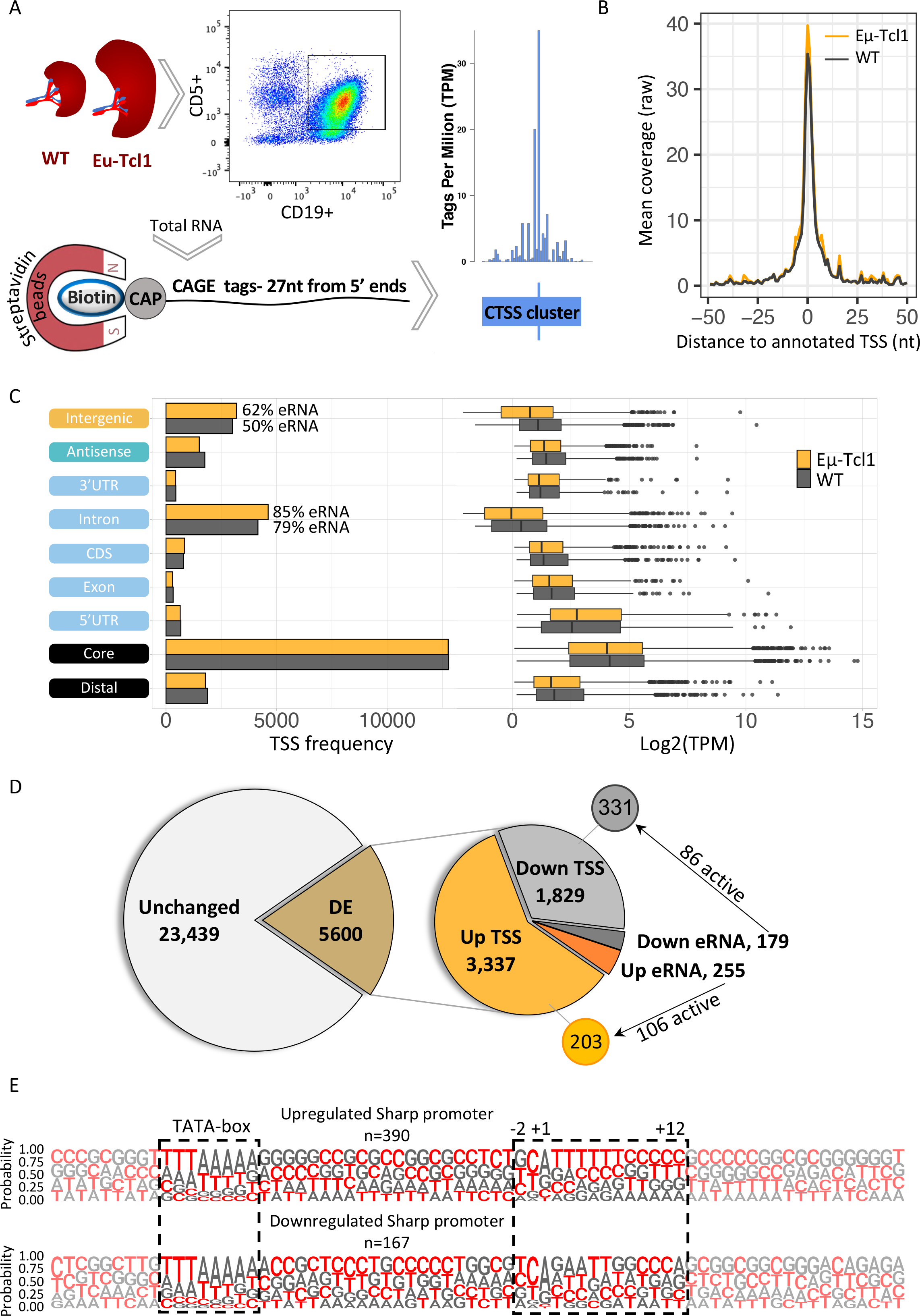
Global analysis of TSSs in healthy and CLL B cells. **A.** A scheme of the Cap Analysis Gene Expression (CAGE) in healthy (WT) and CLL B cells (Eµ-Tcl1). WT and Eµ-Tcl1 splenic B-cells were isolated using CD5 and CD19 magnetic beads. Total RNA samples were subjected to the CAGE as described in the methods section. CAGE-derived Transcription Start Sites (CTSSs) were mapped and clustered into genomic blocks referred as tag clusters (blue bar) with a peak of most recovered CTSS (blue tick) indicates the TSSs in a single base resolution. **B.** Metagene analysis of Eµ-Tcl1 (orange) and WT (grey) CAGE libraries aligned to annotated TSS atlas of the FANTOM5 project. **C.** The frequency (left) and expression level (right, log2 TPM) of CAGE tag clusters in Eµ-Tcl1 (orange) and WT (grey) mice by gene locations. The percentage values on the columns of intronic and intergenic locations refer to predicted eRNA TSSs. **D.** Differentially expressed (DE) TSSs. Left circle indicates unchanged (light grey) and significantly DE (brown) TSS between Eµ-Tcl1 and WT mice. Right circle details frequencies of Eµ-Tcl1 upregulated genic and eRNA TSSs (orange and dark orange, respectively) and downregulated genic and eRNA TSSs (grey and dark grey). Subgrouping of DE TSSs that are positively correlated with enhancers are indicated. **E.** Sequence LOGOs (- 40 to +30 relative to the TSS) of Eµ-Tcl1 upregulated (lower panel) and downregulated (upper panel) sharp promoters where TATA-box situated between -31 to -24 sites and cap proximal region (12 bases) are boxed in a dashed line.

### A marked elevation of new CLL-specific protein isoforms generated by intragenic TSSs

When an alternative TSS is located at the promoter or at the 5’UTR regions, the original ORF is retained. However, new intragenic TSSs located in introns or CDS are predicted to give rise to protein isoforms bearing alternative or truncated N-termini and, consequently, potential different or even opposite biological functions (illustrated scheme in Fig. 2A). We analyzed alternative promoter usage in our data and found that 2,530 genes (22%) contain more than one promoter that contributes over 10% of their total gene expression (Fig. 2B), consistent with previous studies (21). To determine differential TSS usage (DTU), we examined the multi-promoter subset and found 550 significant DTUs from 489 genes where the alternative TSS is upregulated in Eµ-Tcl1 and 373 DTUs from 328 genes with downregulated alternative TSS. The 47% more cases of upregulated alternative TSS indicate that DTU during CLL transformation is gaining more new isoforms than losing canonical ones (Fig. 2C). The Eµ-Tcl1 upregulated alternative TSSs were enriched at the core promoter, 5’UTR, CDS, and intron regions but not at distal regions (Fig. 2C). Median expression (log2TPM) of alternative TSS in all locations ranged between 1.6 to 2.7 with no apparent differences between upregulated and downregulated alternative TSS (Figure 2 – figure supplement 1A). By analyzing the correlation between alternative TSSs and nearby active enhancers, which are marked by eRNAs, a clear elevation in eRNA frequencies was found in Eµ-Tcl1 upregulated alternative TSSs (Figure 2 – figure supplement 1B). These findings suggest the significant role of active enhancers in the activation of DTUs in Eµ-Tcl1.

**Figure 2:**
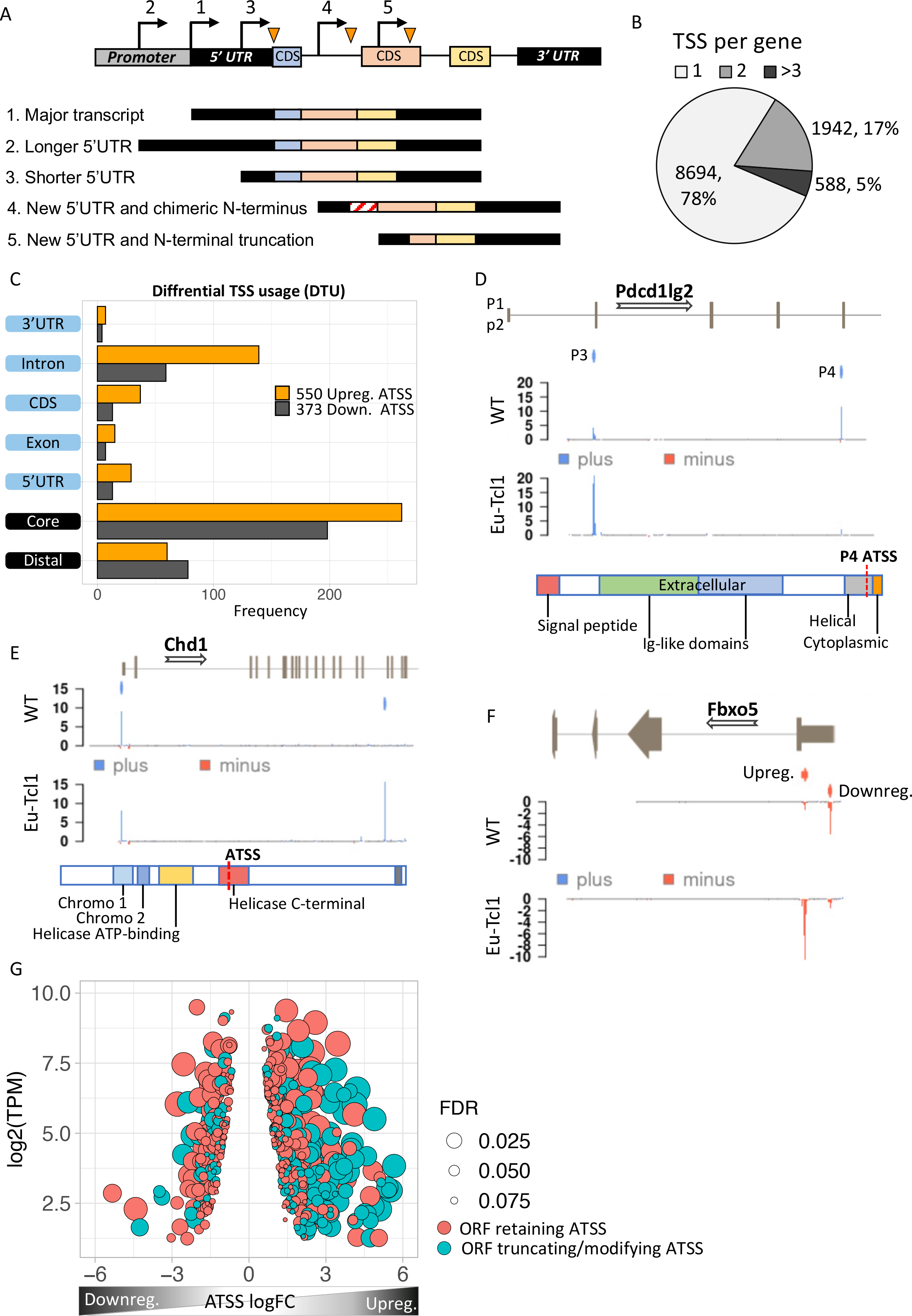
Analysis of differential TSS usage (DTU) reveals alternative 5’ UTRs and new protein isoforms derived from cryptic promoters in CLL. **A.** Scheme of five alternative TSSs (ATSSs) generating isoforms with alternative 5’ UTR regions (1–5) and modification of the N’ termini e.g., new chimeric CDS (3) and N’-terminal truncation (5). **B.** The frequency and the percentage of uni-TSS genes and multi-TSS genes including only TSS clusters that constitutes over 10% of total gene expression. **C.** Frequencies of alternative TSSs that were upregulated (orange) and downregulated (grey) in Eµ-Tcl1, grouped by gene-structure locations. **D.** Gene track showing differential TSS usage (DTU) in Pdcd1lg2 (PD-L2) in Eµ- Tcl1 and WT mice. Canonical promoter (P3) and intragenic promoter (P4, located just upstrem to the last CDS of PD-L2) are upregulated and downregulated in Eµ-Tcl1, respecively. Both, located on the plus strand (blue). Bellow, is a scheme illustrating the P4-induced truncation of PD-L2 lossing most of its functional domains. **E.** Chd1 protein truncation in Eµ-Tcl1 resulted by induction of cryptic promoter. In the upper track, two promoters on the plus strand (blue ticks) presented along to a partial gene-structure scheme. While the canonical promoter located upstream to the annotated 5’ UTR is similarly expressed in Eµ-Tcl1 and WT, an Eµ- Tcl1-specific induction of an intragenic alternative TSS located in the 16^th^ intron resulting in N’ terminus truncation of Chd1 lacking several active domains as illustrated below. **F.** Fbxo5 5’ UTR shorthening resulted by DTU in Eµ-Tcl1. On the upper track, two alternative promoters located on the minus strand shown as short red bars with indicator tick of TSS. Coverage peaks in the two lower tracks (WT and Eµ-Tcl1) showing differential expression of the two promoters where the nearest alternative TSS to the ORF is upregulated in Eµ-Tcl1. **G.** Dot- plot of ORF retaining alternative TSS (ATSS) in peach and ORF truncating intragenic alternative TSS (turquoise) presenting expression (log2 TPM) against ATSS logFC evaluating the degree of promoter shifting (DTU). Positive and negative ATSS logFCs refer to Eµ-Tcl1 upregulated and downregulated ATSS, respectively. Dot sizes correspond to the FDR statistical significance of the DTU analysis.

To explore the consequences of DTUs in CLL, we first analyzed several DTU events (Fig. 2D-F). An intriguing example of DTU resulting in an N-terminally truncated isoform was found in programmed cell death 1 ligand 2 (Pdcd1lg2/PD-L2). This gene has an intragenic alternative TSS generating a protein isoform lacking the Ig-like extracellular domains of PD- L2 (30–32), specifically expressed in WT B cells and is predicted to be inactive. In contrast, the full-length protein is exclusively expressed in the CLL cells via promoter shift and likely contributes to immunosuppression (Fig. 2D). An opposite example is chromodomain 1 (Chd1), in which promoter shifting in Eµ-Tcl1 is within an intragenic alternative TSS that results in truncation of several critical active domains, presumably leading to loss of function (Fig. 2E). The F-box only protein 5 has two promoters located upstream to the main ORF differing only by their 5’ UTR length resulting in 490 nt shorter 5’ UTR in the Eµ-Tcl1-specific transcript isoform (Fig. 2F). Global analysis of the promoter shifts levels and its significance of all DTUs revealed that the upregulated alternative TSS isoforms predicted to give rise to proteins with alternative/truncated N-termini are dramatically increased in the Eµ-Tcl1 samples compared to WT B cells (Fig. 2G). This finding suggests a previously unappreciated diversity of the CLL proteome.

Gene ontology (GO) analysis of genes affected by AP usage shows enrichment in critical CLL pathways, e.g., NF-kappaB signaling (13), MAPK signaling (33), Toll-like Receptor signaling (34), Phosphatidylinositol 3-Kinase signaling (35), Chromatin regulation/acetylation and other relevant GOs (complete list in Figure 2- supplement table 1). Figure 2- supplement table 2 lists the phenotypes derived from mouse genome informatics (MGI) associated with the DTU gene subset, such as Increased B cell number, Enlarged spleen, Thymus hypoplasia, etc, consistent with hematopoietic malignancy.

**Table 1.** Differential expressed chromatin-regulating genes

| **Symbol** | **logFC** | **adj.P.Val** | **Protein** |
| --- | --- | --- | --- |
| Chd2 | -1.28 | 0.04 | Chromodomain helicase DNA binding protein 2 |
| Chd3 | -8.29 | 0.01 | Chromodomain helicase DNA binding protein 3 |
| Cbx4 | -1.69 | 0.02 | Chromobox 4, E3 SUMO ligase |
| Kmt2c | -1.2 | 0.03 | Lysine methyltransferase 2C |
| Kmt5b | -1.47 | 0.03 | Lysine methyltransferase 5B |
| Hdac11 | -13.79 | 0.02 | Histone deacetylase 11 |
| Hdac9 | -4.59 | 0.02 | Histone deacetylase 9 |
| Kat6a | -1.04 | 0.03 | Lysine acetyltransferase 6A |
| Kdm7a | -1.27 | 0.02 | Lysine demethylase 7A |
| Smarca2 | -2.13 | 0.03 | SWI/SNF related |
| Asf1b | 4.36 | 0.032 | anti-silencing function 1B histone chaperone |

### Tcl1 promotes chromatin relaxation, induction of intragenic cryptic promoters, and activation of the c-Myc oncogene

To elucidate the underlying basis for the broad induction of intra-genic cryptic promoters, we considered previous genetic and molecular studies reporting transcription initiation within coding regions as a consequence of defects in the activities of chromatin regulators that maintain closed chromatin structure or nucleosome positioning (36–41). Our Eµ-Tcl1 data found 34 chromatin remodelers with ORF-retaining or ORF-truncating alternative TSSs that could be classified into factors associated with open or close chromatin states, 28 of them were upregulated in Eµ-Tcl1 (Fig. 3A). Remarkably, many of the upregulated epigenetic gatekeepers of coding regions such as Hdac4, Hdac5, Sirt2, Smyd4, Kdm4a, Dnmt3a, Chd1 and Mgmt, are predicted to express truncated proteins that cause loss of critical domains due to DTU (Fig. 3A and 5D). Moreover, numerous other epigenetic regulators of closed chromatin structure are strongly downregulated (Table 1). These findings revealed widespread chromatin deregulation in Eµ-Tcl1 B cells that can activate cryptic promoters.

**Figure 3:**
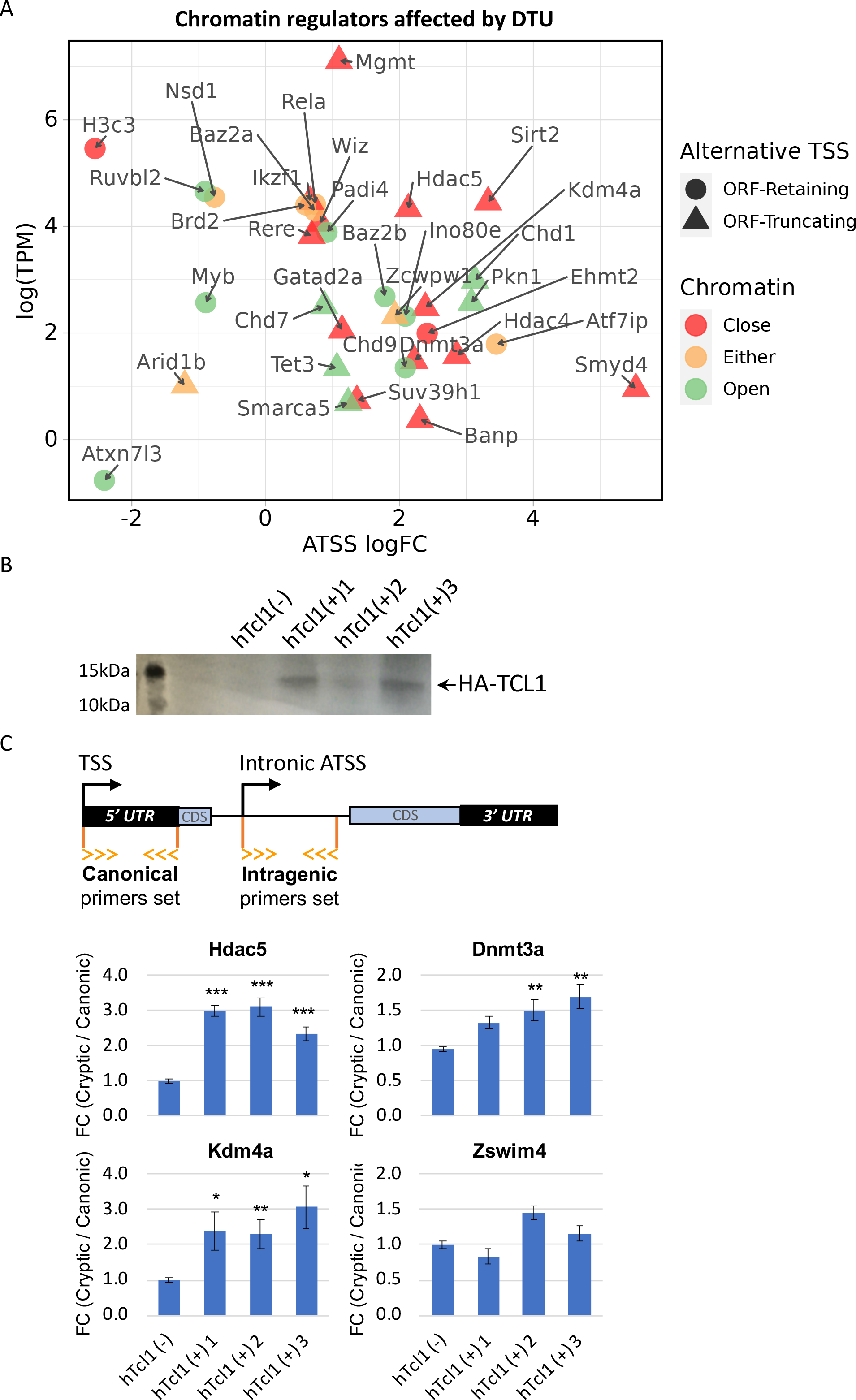
Cryptic promoters of closed chromatin epigenetic regulators and the role of hTcl1. **A.** Dot-plot of ORF-retaining (circle) and ORF-truncating (triangular) alternative TSSs within genes associated with close (red), open (green), or either (yellow) chromatin states. For evaluating the degree of promoter shifting (DTU), the data is presented by expression (log2 TPM) against logFC of alternative TSS (ATSS). Positive and negative ATSS logFCs refer to Eµ-Tcl1 upregulated and downregulated ATSS, respectively. **B.** Generation of hTCL1 expressing MEFs. MEFs cells were transfected with hTCL1 expression plasmid, co-expressing the Neomycin gene and 3 stable clones expressing HA-tagged hTCL1 were selected using G418 antibiotics. TCL1 expression was verified by western blot with anti-HA antibody. **C.** Exogenous expression of hTCL1 induces cryptic promoters. A scheme illustrating the canonical and cryptic promoter and the design of RT-qPCR primers to the 5’ UTR sequences originated by canonical TSS and intragenic (intronic) alternative TSS. The results are presented as the ratio between cryptic and canonical levels. Statistically significant differences are denoted by asterisks as follows: *P*< 0.05= *, ** = *P*< 0.01 and *** = *P*< 0.001).

The induction of intra-genic promoters can be either driven by Tcl1 (intrinsic) or by environmental signals associated with cancer progression. To distinguish between these possibilities, we established three mouse embryonic fibroblast (MEFs) clones stably expressing human Tcl1, and determined, by RT-qPCR, the relative expression of the alternative intra-genic TSS of 4 selected genes (as shown in the scheme). We observed a significant elevation of the intra-genic truncating TSS of Hdac5, Dnmt3a, and Kdm4 but not Zswim4. These findings suggest that Tcl1 itself can promote relaxation of the chromatin structure and promoter shifts, at least in some genes.

De-regulation of the chromatin affects the accessibility of TFs in promoter and enhancer regions. Since the CAGE provides expression data of genes along with their promoter and enhancer regions, we investigated the contribution of TFs to the activation of cryptic promoters in Eµ-Tcl1. By generating position frequency matrices (PMSF) of TFBS motifs (JASPAR 2020, mouse core collection) we could appreciate the apparent differences in the occurrence of TFBS in ORF-retaining and ORF-truncating alternative TSSs, differentially transcribed in Eµ-Tcl1 (Fig. 4A). Shown by PCA2, TFBS frequencies mainly differ between upregulated and downregulated ORF-truncating alternative TSS, explaining 38.6% of the variation (Fig. 4A). Using Fisher’s exact test, we found 63 TFBS enriched in the promoters of the upregulated alternative TSSs compared to the promoters of the canonical upregulated TSS in Eµ-Tcl1 (Fig. 4B). When we combine this data with the list of up-regulated TFs in Eµ-Tcl1 (Table S3), we found that upregulated levels of the proto-oncogenes Ets2 and Myc correspond to the highly enriched Ets-related and bHLH motifs, respectively (Fig. 4B).

**Figure 4:**
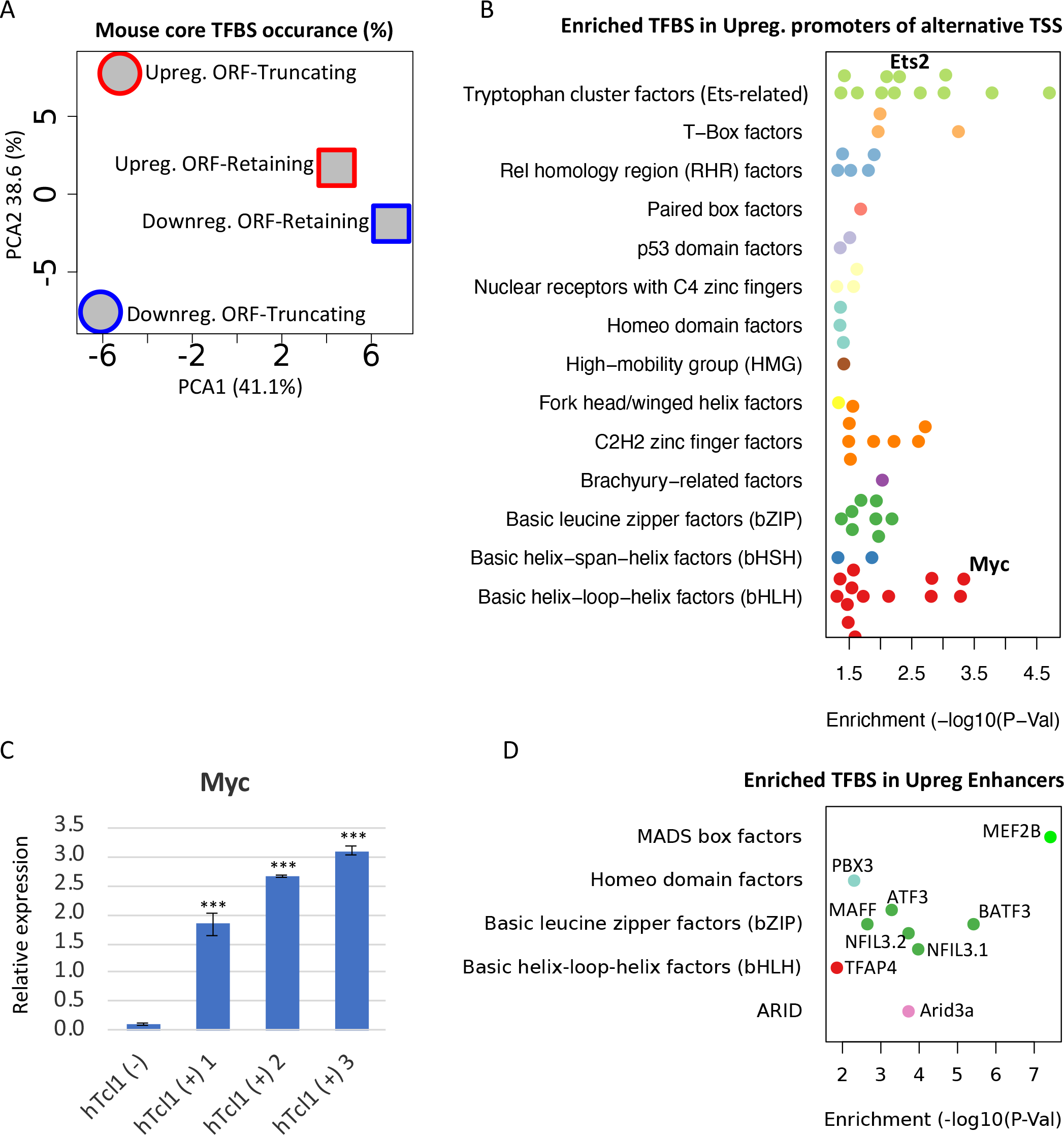
Upregulation of TFs targeting cryptic promoters and enhancers. **A.** A PCA analysis of motif occurance (%) of 119 transcription factor binding sites (JASPAR 2020, mouse core collection) searched in 923 promoters (-1000 upstream and +100 downstream to TSS) of upregulated (red) and downregulated (blue), ORF-retaining (squares) or ORF-truncating (circles) alternative TSSs. PCA1 and PCA2 explain 38.6% and 41.1% of the total variation, respectively. **B.** TFBS enrichment analysis of activated alternative promoters (-1000 upstream and +100 downstream to TSS) relative to the overall activated promoters. Among the enriched TFs, Ets2 and c-Myc levels are upregulated in Eµ-Tcl1 (Table S3) and are indicated. **C.** Analysis of the c-Myc expression in the stable MEF clones expresing hTcl1 by RT-qPCR. Statistically significant differences are marked with asterisks *** = *P*< 0.001. **D.** TFBS enrichment analysis of enhancers associated with alternative promoters relative to overall induced enhancers. The levels of indicated enriched TFs are upregulated in Eµ-Tcl1 (see Table S3).

Activation of the c-Myc oncogene is the hallmark of many hematopoietic malignancies (42). As c-Myc is among the upregulated TFs in our CAGE data of Eµ-Tcl1 CLL and is predicted to regulate intra-genic alternative promoters, we asked whether Tcl1 itself can activate c-Myc. Using the hTcl1 MEFs clones, we found that the levels c-Myc are dramatically up-regulated in all the hTcl1 expressing clones, indicating that its activation in Eµ-Tcl1 CLL B is most likely intrinsic.

As many of the upregulated alternative TSSs in Eµ-Tcl1 are associated with active enhancers (Figure 2 – figure supplement 1B), we analyzed TFBS enrichment also in the upregulated enhancers and found nine enriched motifs (Fig. 4D). Remarkably, the TFs linked to these nine enriched motifs are all upregulated in the Eµ-Tcl1 (Figure 4 – supplement table 1) and include MEF2B, PBX3, MAFF, NFIL3, ATF3, BATF3, TFAP4, and ARID3A. These findings strongly suggest that these enhancers and their corresponding upregulated transcription factors act as inducers of DTUs in Eµ-Tcl1. Altogether, these findings expand the oncogenic activities of hTcl1 to de-regulation of diverse chromatin factors, activation of the c-Myc oncogene and enhancer-specific TFs, which cooperate to induce intra-genic cryptic promoters.

### Polysome-CAGE analysis of DTU in Eµ-Tcl1 CLL B cells reveals high translatability of intragenic (cryptic) APs and impairment of epigenetic regulation

Considering that the extensive changes in TSSs are predicted to generate transcript isoforms that vary in their 5’UTR and translation start site, we examined whether the Eµ-Tcl1-specific mRNAs originated from DTUs are translated into proteins. To this end, we performed polysome profiling of Eµ-Tcl1 splenic B cells and generated CAGE libraries from the fractions representing polysome-free (Free), light polysomes occupying two to four ribosomes (Light), and heavy polysomes (Heavy) occupying over four ribosomes per mRNA transcript (Fig. 5A). The polysome profile of the Eµ-Tcl1 CLL cells shows a high monosome to polysome ratio (Fig. 5A), reminiscent of a state of translation initiation halt, suggesting that protein synthesis capacity is limited in these cells. 3.39 million CTSSs were mapped and clustered to a consensus set of 30,246 promoters corresponding to 11,465 genes. Similar to the total RNA CAGE samples, most TSSs of the polysome profiling samples mapped with the highest expression level to the core promoter region as opposed to the highly frequent but poorly expressed intergenic and intronic TSSs (Fig S3A). As a support for the eRNA prediction shown to be abundant in the intergenic and the intronic regions (Fig. 1C), most intergenic and intronic TSSs were found in the polysome-free fraction (Figure 5 – figure supplement 1A and B), confirming that those entities are non-coding RNAs transcribed from active enhancers. To determine the translation efficiency (TE) of isoforms generated by alternative TSSs of multi-TSS genes, we calculated the ratio between the TSS counts in Heavy + Light fractions and the Free fraction. To explore the consequences of alternative TSS on translation efficiency, we defined “Differential High-TE” only if its TE value was 2-fold higher than its paired TSS, and vice versa for “Differential Low-TE” alternative TSS (illustrated in Figure 5 – figure supplement 1C). Out of 13,924 TSS pairs, 2,458 were assigned as “Differential High-TE” and 2,103 classified as “Differential Low-TE”. After extracting 5’UTR sequences (see Material and Methods), we tested our High/Low-TE calling by evaluating known translation regulatory features of 5’UTR. We found that “Differential Low-TE” TSSs tend to have longer 5’UTR and to contain more uORF (Fig. 5B), which are well-known inhibitory translation features.

**Figure 5:**
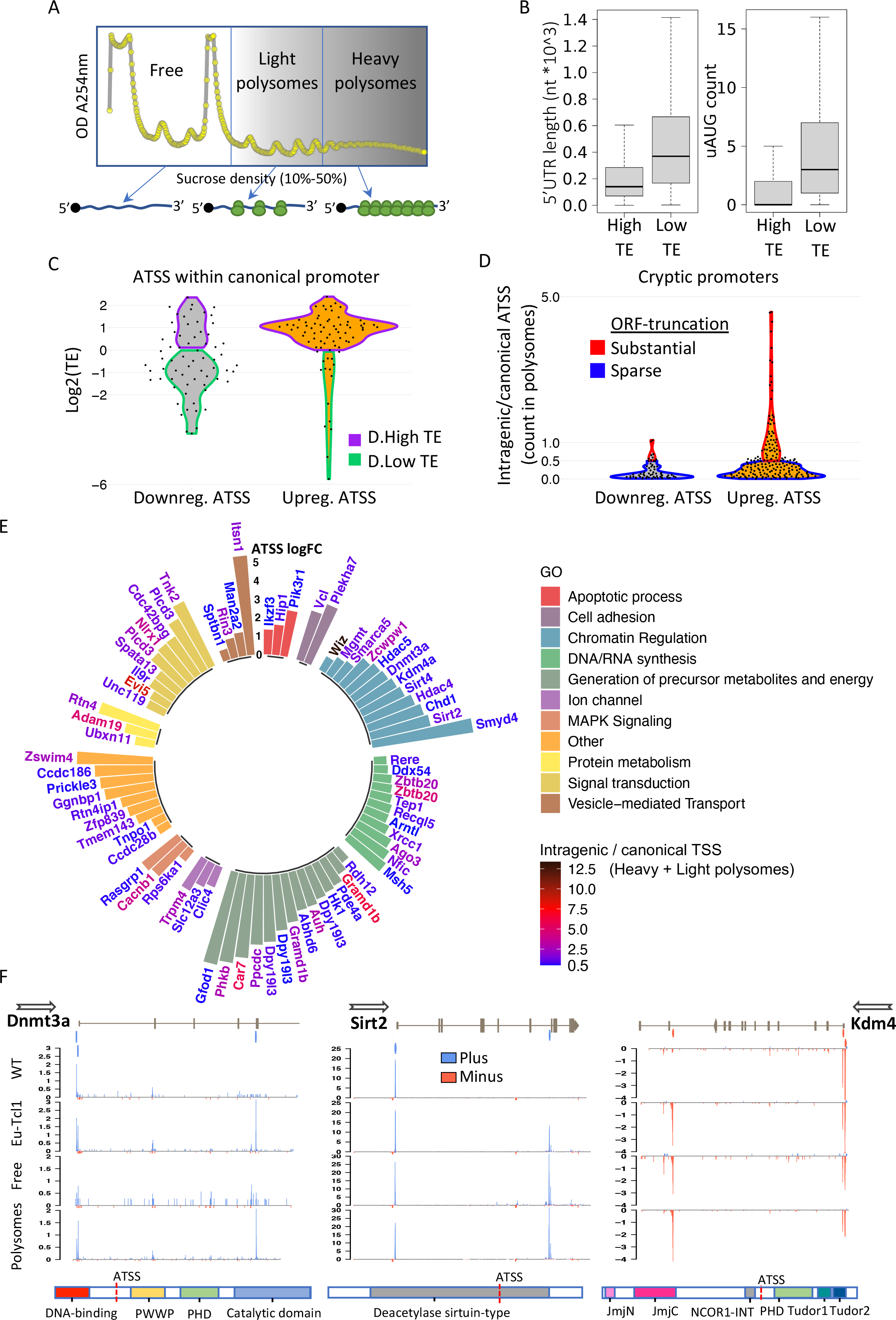
Translatability of ORF-retaining and ORF-truncating alternative TSSs. **A.** Schematic presentation of polysome fractionation into three main fractions. Free fraction corresponds to non-translated transcripts and Light and the Heavy fractions corresponds to intermediate and highly translated transcripts, respectively. **B.** Boxplot of 5’ UTR length and number of upstream AUG of transcript isoforms defined by high translation efficiency (High-TE) and low translation efficiency (Low-TE). **C.** The TE distribution of differentially expressed ORF-retaining alternative TSSs (within canonical promoter) leading to differential High-TE (purple border) and differential Low-TE (green border) transcript isoforms. **D.** ORF-truncating ATSSs that are either above (Substantial ORF truncation) or below (Sparse ORF truncation) 50% of the canonical TSS count in the polysomes fraction. **E.** Gene Ontology (GO) anlysis of genes with Eµ-Tcl1 upregulated and translated intragenic ORF-truncating ATSSs. Bar hights presents an inclusive logFC explaining the magnitude of promoter shifting, taking in consideration TSS FC between Eµ-Tcl1 and WT mice and FC between cannonical and intragenic TSSs paires, per gene. Gene names are colored by a relative scale of translation level between ORF-truncating ATSS and the canonical TSS, calculated by the ratio of intragenic / canonical TSS counts in the polysome fractions. Blue colored gene reffer to ORF-truncating ATSS that exceeds 50% (0.5) of the canonical representation in the polysomes fraction and dark red reffer to the maximum of 12.5 FC higher representation in the polysomes fraction by the ORF-truncating ATSS. **F.** Coverage peaks of TSSs of canonical and intragenic cryptic promoters (indicated by blue and red ticks in the upper track) found in Dnmt3a Sirt2 and Kdm4 genes. Blue and red coverage peaks represents transcription from the plus and minus strands, respectively. Eµ- Tcl1 and WT tracks presented TSS peaks of total RNA and Polysomes and Free tracks derived from Eµ-Tcl1 polysome profilling. On the bottom are schemes of each gene’s protein domains in which the points of truncation due to intragenic ATSS are marked.

Next, we examined the translatability of Eµ-Tcl1-specific transcript isoforms that arise from promoters shifts (DTU). By intersecting the CAGE data of total RNA with polysome profiling fractions, we could determine the translatability of each isoform initiated by alternative TSS. Out of 543 DTUs found within the 5’ UTR, 142 were also differentially translated, of which 76 and 66 were transcriptionally up- and down-regulated, respectively. Intriguingly, the vast majority of the transcriptionally upregulated DTU (60/76) are classified as High TE, and most of the downregulated DTUs display Low TE (41/66) (Fig. 5C), indicating for coordination of TSS selection with mRNA translation.

To assess the potential functional consequences of the intragenic alternative TSSs predicted to generate N-terminally modified protein isoforms, we compared the polysomal CTSS tags of the truncated isoform produced by the intragenic alternative TSS to the polysomal level of the full-length isoform produced by the canonical TSS. When the intragenic alternative TSS counts in the Heavy and Light polysomal fractions exceed 50% of the canonical TSS, we consider this cryptic transcription to potentially affect the function of the annotated full-length protein. In total, we found 73 intragenic alternative TSS that are upregulated in Eµ-Tcl1 and are translated ≥ 50% of the canonical transcripts and 168, which their lower translatability will less likely to impact protein functionality (Fig. 5D). GO analysis of these 73 genes is summarized in Fig. 5E by plotting the level of promoter shifting (scored as inclusive logFC) and the relative translation level of an intragenic alternative TSS and canonical TSS, which provides insight into the potential vulnerability of cellular functionalities. Genes have been clustered into the following cellular processes: ‘apoptotic process’, ‘cell adhesion’, ‘chromatin regulation’, ‘DNA/RNA synthesis’, ‘generation of precursor metabolites and energy’, ‘ion channel’, ‘MAPK signaling’, ‘protein metabolism’, ‘signal transduction’ and ‘vesicle-mediated transport’. Interestingly, under ’chromatin regulation’, we could find Eµ-Tcl1- specific truncated isoforms of chromatin remodelers and epigenetic players that might be involved in the aberrant TSS utilization found in Eµ-Tcl1 B cells. This class includes histone modifiers: Smyd4, Sirt2, Sirt4, Hdac4, Hdac5, Kdm4a, Zcwpw1, and Wiz; DNA methylation factors: Dnmt3a and Mgmt and chromatin remodeling factors: Chd1, Smarca5, and Bcl7b. In addition, under ’DNA/RNA synthesis’ GO, we could find Eµ-Tcl1-specific truncated isoforms of DNA repair genes such as Xrcc1, Recql5, and Tep1, and direct transcription regulators essential for hematopoiesis that includes Gfi1, Zbtb20, Ago3, Ssbp4, Ddx54, and Arntl (the complete list with descriptions detailed in Table S3). These cases of promoter shifting in Eµ- Tcl1 leads to the generation of highly translated protein isoforms with possible loss of function or dominate-negative activities. For example, Eµ-Tcl1-specific cryptic transcription (2.22 logFC) within the fourth intron of Dnmt3a leads to the loss of three exons (211 amino acids) that constitute its DNA-binding domain (Fig. 5F), which is important to the DNA methylation activity (43) and is generally associated with closed chromatin state(36). The Dnmt3a cryptic alternative TSS is enriched within Polysomes fractions, suggesting that Dnmt3a activity is substantially impaired (Fig. 5F, left). A promoter shifting in Sirtuin 2 (Sirt2, 3.32 logFC), a type III histone deacetylase (HDAC), interrupts the deacetylase domain positioned between residues 65 and 340 that generates a truncated isoform with higher translatability compared to the canonical isoform and is enriched in the polysomes fractions (Fig. 5F. middle). Together with HDACs, lysin demethylase (KDMs), erasers of histone three lysin-9 and lysin-36 methylation marks of active transcription are known to be co-recruited to ensure gene silencing(38). DTU found in lysine-specific demethylase 4a (Kdm4a) has an Eµ-Tcl1-specific cryptic transcription leading to truncated isoform that is equally presented in polysome fractions as the canonical full-length isoform (Fig. 5F, right). Thus, with five truncated members of the HDAC and KDM families (Hdac4, Hdac5, Sirt2 and Sirt4, Kdm4a) and dramatic downregulation of Hdac11 and Hdac9 (-13.79 and -4.59 log FC, respectively, Table 1), it appears that Eµ-Tcl1 B cells have reduced eraser activities of histone acetylation and methylation, to maintain open chromatin state.

### Identification of CNY initiating trinucleotides as translation inhibitory motif associated with metabolic stress signature

The polysome-CAGE data enable the determination of the exact first nucleotide of every transcript and its impact on translation efficiency. We first determined the frequencies of TSS nucleotides in our Eµ-Tcl1 data and found that the frequency of A and G (47% and 36%, respectively) is higher than the C and T (9% and 4%, respectively), as expected (Fig. 6A). Yet, the difference between purines to pyrimidines is substantially more significant. For example, in Eµ-Tcl1, purines account for 83%, but in MEFs, only 64% (21), suggesting an increased preference of purines as start sites in B CLL cells. We next determined the TE of transcripts with different initiating nucleotides and observed that those initiated with a “C” have a substantially lower TE compared to the A, G, and T (Fig. 6B). To examine whether the lower TE associated with the cap-proximal C is linked to the well-known TOP element (CYYYY), we analyzed the TE of the first trinucleotides and found that almost all C initiating trinucleotides have a relatively lower TE, including inhibitory trinucleotides that are part of the TOP element such as CCT and CTT, but also others that deviate from the TOP such as CAC, CAT and CTG. Interestingly, among the C-initiating triplets, those with a pyrimidine (CNY) in the third position are associated with a much lower TE (Fig. 6C). Importantly, the nucleotides that follow the CNY do not seem to contribute to the TE suppression (Fig. 6D), indicating that the CNY context is the predominant feature. Fig. 6E presents three examples of TSSs adjacent to each other within the same promoter region and displays a differential polysome to free ratio. In the Mdh2 gene, the reads of the two C in the CAC triplet are more enriched in the Free than the Polysomal fractions, while the opposite is seen for the reads that start with the A. The same trends are also seen in Psmb4 and Atp5f1, of which the As are more highly translated than the Cs. As the 5’UTR generated from these TSSs is almost identical, these examples highlight the important contribution of the first nucleotides to translation efficiency.

**Figure 6:**
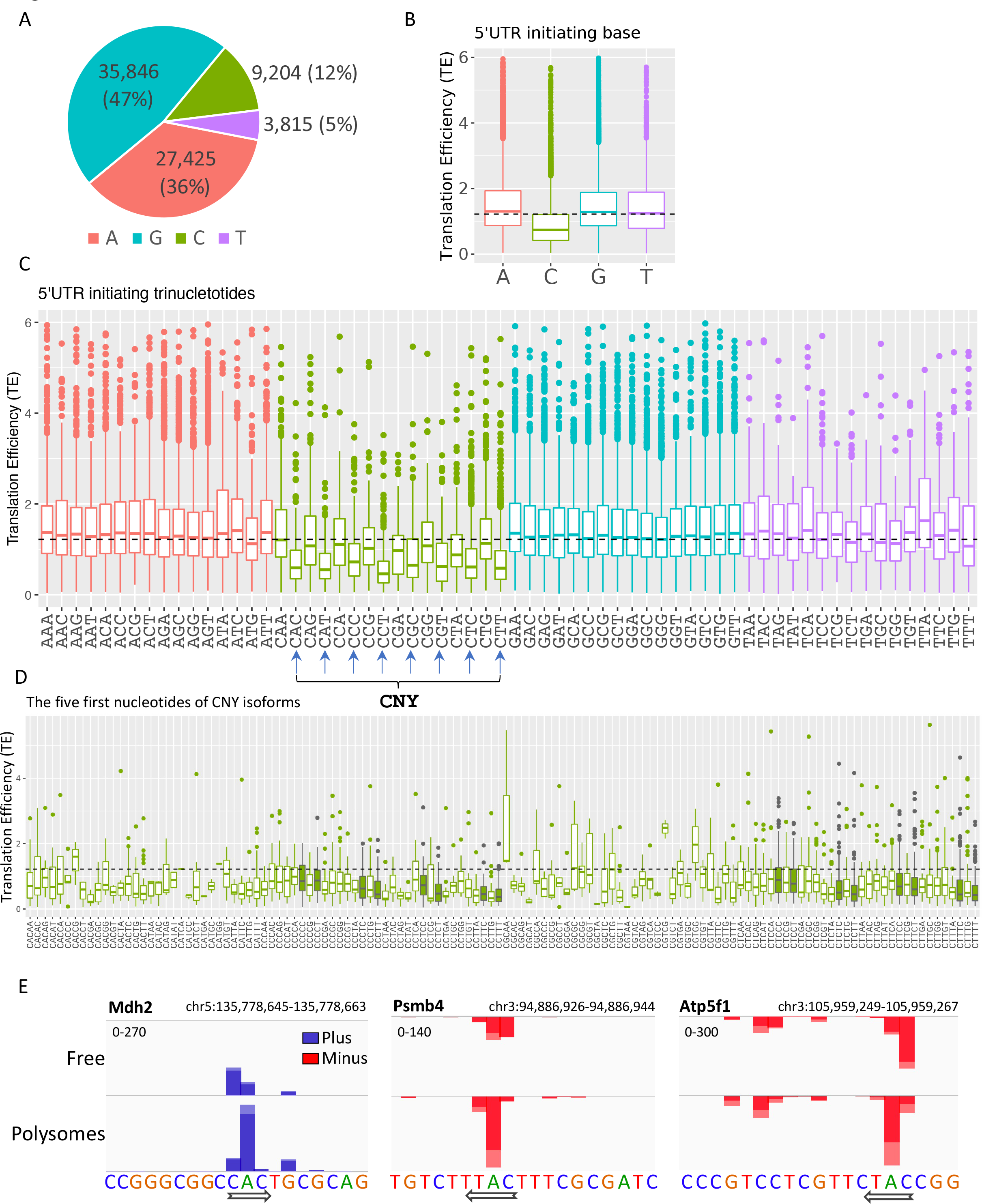
The impact of TSS nucleotides on translation efficiency. **A.** The frequency of initiating TSS nucleotide in Eµ-Tcl1. **B.** Boxplots of translation efficiency (TE) of isoforms that differ in their TSS first nucleotide. The horizontal dashed line marks the overall median value. **C.** Boxplots of TE of transcripts that differ in their first three nucleotides. CNY trinucleotides are marked by arrows. **D.** Boxplots of TE of the first five nucleotides of the CNY-initiating transcripts. The nucleotides corresponding to the TOP element are marked with filled boxes. All the data presented in this figure are the mean of the two independent replicates. The bottom and the top whiskers represent 5% and 95% of the distribution, respectively. **E.** Examples representing the effect of the first nucleotides on translation efficiency in Mdh2, Psmb4 and Atp5f1 genes. CTSS tags per million (TPM) is plotted separately for plus (Blue) and minus (red) strands of the polysome-free and polysomal fractions with a scale set for the two tracks shown in the upper-left corner. Direction of transcription is marked by an arrow located at the first CTSS.

A previous study that analyzed the relationship between the first nucleotide and the relative translation efficiency also reported that “C” initiating transcripts are associated with lower TE, particularly following energy stress (21). We re-analyzed this previous data and found that in energy-starved MEFs, the CNY is also associated with more significant inhibition of the TE (Figure 6 – figure supplement 1A and B). Thus, the higher monosome to polysome ratio (Fig. 5A) and the association of “CNY” with reduced TE are consistent with Eµ-Tcl1 B CLL cells being under metabolic stress.

### CLL B cells are adapted to the decline in epigenetic, DNA repair, and translation activities

To examine how CLL cells cope with the limited activities of epigenetic regulators, DNA repair, and translation initiation, we isolated and cultured CLL and non-CLL B cells from Eµ-Tcl1 mice as well as B cells from healthy mice and treated them with drugs specific to these pathways. Specifically, we used inhibitors against the downregulated epigenetic factors HDAC3A, HDAC2A, KDM4A, and the rate-limiting translation initiation factor eIF4E, in a dose-response assay. We found that the non-CLL or healthy B cells were inhibited by the HDAC2A and eIF4E inhibitors in a dose-dependent manner, whereas the CLL cells displayed either resistance (eIF4Ei) or reduced sensitivity (HDAC2Ai) (Fig. 7). Interestingly, none of the B cell types was affected by inhibitors against HDAC3A and KDM4A at the concentrations used. These findings suggest that the CLL cells are substantially less dependent on the activities of these limiting factors and are well adapted to their reduced activity.

**Figure 7:**
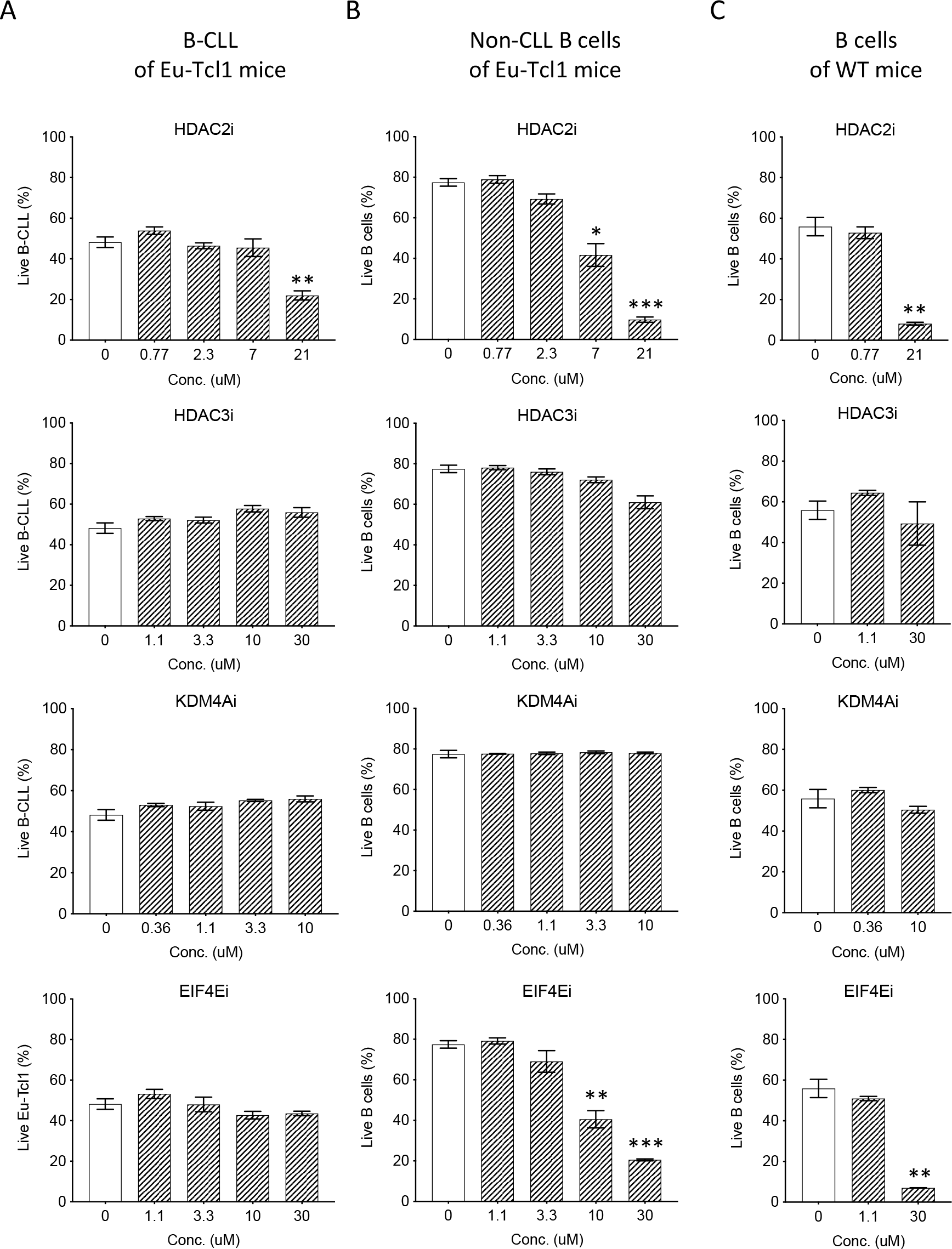
Eu-Tcl1 B cells are relativlely resistant to inhibitors of affected pathway. **A-C**. Survival assay of B-CLL of Eu-Tcl1 mice (A), non-CLL B cells of Eu-Tcl1 mice (B), B cell of WT mice (C) treated with inhibitors (i) targeting HDAC2, HDAC3, KDM4A and EIF4E. The significance between DMSO (vehicle control) and inhibitor-treated cells is indicated by asterisks (* for *P*<0.05, ** for *P*<0.01 and *** for *P*<0.001).

## Discussion

In this study, we used the Eµ-Tcl1 mouse model of Chronic Lymphocytic Leukemia to investigate the global changes in TSSs and AP usage and their impact on the protein content of cells undergoing oncogenic transformation. By comparing the TSSs landscape of normal and CLL B cells, our findings revealed a dramatic induction of APs in malignant CLL B cells with a particularly high prevalence of intragenic promoters predicted to generate N-terminally truncated or N-terminally modified proteins. Importantly, a major class of intragenic alternative promoters activated in the Eµ-Tcl1 CLL cells is the ’closed chromatin’ epigenetic regulators gene subset, predicted to generate a loss of function or dominant-negative isoforms. Enhanced chromatin accessibility due to down-regulation of epigenetic gatekeepers is well-known to promote transcription initiation from intra-genic cryptic promoters sites (36–41). Thus, intragenic promoter shifts in B CLL follow a feed-forward loop, in which activation of N-terminally truncated closed chromatin modifiers further enhance the cryptic promoter activity. Additionally, the regulatory regions of the intragenic-cryptic initiation sites (proximal promoters and the nearby enhancers) are targeted by the c-Myc oncogene and several transcription factors that are specifically upregulated in Eµ-Tcl1 B CLL. Thus, the combined loss of epigenetic regulators and the induced expression of a specific subset of transcription factors underlie the dramatic TSSs and AP usage changes in Eµ-Tcl1 B CLL. Notably, these dysregulated gene expression features are most likely intrinsic to the Eµ-Tcl1 B CLL, as exogenous expression hTcl1 in MEFs is sufficient to drive the induction of intragenic promoter of epigenetic regulators and to up-regulate c-Myc.

Our findings are consistent with a recent study that reported changes in promoter utilization in multiple cancer types (17). As this study used RNA-seq data to infer promoter activities, this data is substantially less accurate for estimating intragenic promoter activity especially when it is embedded within internal exons or introns. Thus, our CAGE data of healthy and malignant B cells provided further insights on intragenic promoter activities, their impact on the translatome, and the mechanism underlying their activation.

A previous study described decreased methylation levels in Eµ-Tcl1 mice and in CLL patients (15) and reported a strong interaction between TCL1 and *de novo* DNMT3A and 3B along with inhibition of the enzymatic activity, suggesting for inhibition of *de novo* methylation during leukemogenesis. Our findings complement these observations by uncovering another mechanism of inhibition of DNM3A via activation of an intragenic promoter and generation of large amounts of truncated inactive protein (Fig. 5F, left). These two mechanisms are likely to synergize to inhibit the DNA methylation in these cells. Notably, an earlier study provided conflicting results and demonstrated increased methylation levels in human as well as mouse CLL cells (44). These discrepancies may reflect the complex timing of epigenetic changes during the progression of the leukemic process.

The use of Polysome-CAGE enabled us to assess the potential contribution of the N-terminally truncated/modified protein isoforms derived from intragenic promoters to the functionality of the intact protein. This analysis reveals substantial impairments in several regulatory and metabolic pathways, including chromatin regulation and energy metabolism, that are expected to contribute to the transformed phenotype. Remarkably, upon analysis of the relationship of the exact cap-proximal nucleotides and translation efficiency, we observed that all “C” initiating transcripts display lower translation efficiency compared to the A, G, and T. Furthermore, among the “C” initiated transcripts, those that start with CNY display even lower TE relative to the other Cs. The effect of the “C” on the TE is nicely demonstrated in the examples in which adjacent nucleotides from the same promoter region display dramatic differences in polysome occupancy (Fig. 6E). In these instances, the 5’UTR length and sequence are almost identical. Considering that translation downregulation of the well-known C-initiating TOP element is the hallmark of various metabolic stresses and mTOR inhibition (45), and the previous observation of downregulation of all C-initiating nucleotides following energy stress (21), we infer that the Eµ-Tcl1 cells display a clear metabolic stress signature. Furthermore, the re-analysis of the first trinucleotide context of the energy-starved MEFs also identified CNY as the most inhibitory signal and the resemblance of the Eµ-Tcl1 CLL to metabolically stressed cells. Thus, our findings expand the regulatory sequences that mediate the effect of stress on translation beyond the TOP element. Such differences in the TE of initiating nucleotides were linked to reduced eIF4E levels upon energy stress, as its cap-binding activity is influenced by the identity of the first nucleotide (21).

Considering the profound effect of TCL1 on multiple major regulatory pathways (Fig. 5) and the adaptation of TCL1 expressing cells to the inactivation of these processes (Fig. 7), it is reasonable that the most effective way to treat these malignancies is to target the TCL1 itself instead of the affected pathways. While TCL1 inhibitors have not yet been discovered, identifying such compounds should be an important therapeutic direction against TCL1-driven malignancies. Notably, the discovery of highly translated N-terminally modified proteins forms the basis for the identification of new, CLL-specific antigens that can be utilized for specific targeting of the CLL cells. Combining inhibitors against TCL1 and immunotherapy could be a promising strategy against CLL.

In summary, the ability of Tcl1 to promote activation of intragenic cryptic promoters of epigenetic regulators results in a feed-forward loop in which the loss of function of ’closed chromatin’ inducers, by expression of N-terminally truncated isoforms, further enhance the activities of the cryptic promoter. This feature of Tcl1 and its ability to boost c-Myc oncogene levels expand the current knowledge on Tcl1 transforming capacities and its multiple modes of regulation. The utilization of the polysome-CAGE not only verified the expression of N-terminally modified protein isoforms but also uncovered the restricted protein synthesis capability of Eµ-Tcl1 B CLL as a point of vulnerability that can be exploited as a drug target. Our findings form the basis for future work on approaches to interfere in tumorigenic processes mediated by TCL1 in TCL1-overexpressing leukemias.

## Methods

### Mice

C57BL/6 wt were purchased from Harlan Biotech Israel (Rehovot, Israel). Transgenic Eμ-TCL- 1 mice (7) were kindly provided by CM Croce (The Ohio State University, Columbus, OH, USA). TCL-1 mice were backcrossed for several generations to C57BL/6 mice to obtain TCL- 1 mice with the same WT background (co-isogenic). Mice were used when they reached progressed illness at the age of one year. All procedures were approved by the Animal Research Committee at the Weizmann Institute.

### Isolation of primary splenic B cells

Healthy and CLL splenic B cells were isolated from one-year-old WT and Eμ–TCL1 mice by positive B-cell selection with CD19 magnetic beads (CD19 MicroBeads, Miltenyi Biotec, cat. 130-121-301). Briefly, spleens were squashed, treated with red blood lysis buffer (homemade) and filtered in PBS using 40-mm cell strainers. The cells were then counted and the B cells were purified according to the manufacturer’s protocol. Splenocytes subjected for polysome profiling were isolated using buffers (above) supplied with 100 ug/ml Cycloheximide (Sigma-Aldrich) to form 80S-mRNA complex.

### Flow cytometry

Analysis of the surface expression of CD5 and CD19 on primary mature WT and Eµ-Tcl1 B cells was monitored by flow cytometry. Briefly, 1 × 10^6 cells were washed with PBS supplemented with 0.5% BSA and 0.1% sodium azide and stained with FITC-conjugated anti- CD5 (eBioscience, cat. 11-0051-85) and with PE-Cy7-conjugated anti-CD19 (Invitrogen , cat. 25-0193-82) antibodies. Stained cells were washed and analyzed by flow cytometry (FACSCanto II, BD Biosciences), and the quantification of the measurements was analyzed using FlowJosoftware (TreeStar). The purity of WT mature B cells was estimated to be between 95% and 99%, and positive-gated Eµ-Tcl1 B cells were estimated between 80% and 85%.

### Polysome profiling

WT and Eµ-Tcl1 B cells were washed with cold buffer containing 20 mM Tris pH 8, 140 mM KCl, 5 mM MgCl2, and 100 ug/ml Cycloheximide. The cells were collected and lysed with 500 ml of the same buffer that also contains 0.5% Triton, 0.5% DOC, 1.5 mM DTT, 150 units RNAse inhibitor (Eurx, cat. E4210), and 5 ul of protease inhibitor (Sigma-Aldrich, cat. 78430). The lyzed samples were vortexed and centrifuged at 12,000g at 4°C for 5 min. The cleared lysates were loaded onto 10–50% sucrose gradient and centrifuged at 38,000 RPM in a SW41 rotor for 105 min at 4°C. Gradients were fractionated, and the optical density at 254 nm was continuously recorded using ISCO absorbance detector UA-6. The collected samples were then pooled to create three main fractions: Polysome-free (Free), Light (2-5 ribosomes), and Heavy (over five ribosomes).

### RNA extraction

Total RNA from WT and Eµ-Tcl1 purified B cells and the RNA from the pooled polysomal fractions (described above) were extracted in two biological replicates using TRIzol reagent (Invitrogen, cat. 15596026) following manufacture’s instructions. After phase separation, the aqueous phase was diluted back in TRizol reagent (1:3) to proceed with RNA purification by zymodirect RNA kit (Zymo research, cat. R2051). The quality of RNA samples was assessed using Agilent 2200 TapeStation (Agilent Technologies, USA) to evaluate RNA integrity (RIN), and purity was measured by Qubit 4 fluorometer (ThermoFisher, cat. Q32852).

### CAGE library preparation and sequencing

Five ug RNA samples were subjected to library preparation of Cap Analysis Gene Expression CAGE using CAGE-seq protocol adapted for Illumina sequencing (23). Specifically, to promote complementary DNA (cDNA) synthesis through GC-rich sequences in the 5’ untranslated regions (5’ UTR), the reaction was carried with the presence of D-trehalose (Sigma-Aldrich cat. T0167) and D-sorbitol (Wako, cat. 19803755) at high temperature. Capped RNAs of RNA-DNA hybrids were Biotinylated (Vector lab, cat. SP1100) and purified by MPG streptavidin beads (Takara, cat. 6124A) followed by RNAse (Promega, cat. M4261) digestion to release cDNAs corresponding to the 5’ ends of the original mRNA. 5’ linkers harboring barcodes sequences and EcoP15I (NEB, cat. R0646S) recognition site were ligated, and second-strand synthesis was performed. Then, fragments of the first 27-base pair (bp) of the 5’ UTR were produced by EcoP15l digestion and ligated to 3’ linkers containing the Illumina primer sequence. The resultant CAGE tags were amplified by polymerase chain reaction (PCR), purified and sequenced by Hiseq 2500 (Illumina) with the addition of 30% PhiX spike-in to balance the low complexity of the 5’ ends of the CAGE libraries.

### CAGE-seq data preprocessing and mapping

Preprocessing pipeline of CAGE raw reads included quality check (QC) using FSTQC, removal of Phix reads (Escherichia virus PhiX174, NC_001422.1) and rRNA CAGE tags using Bowtie2 (2.3.5.1), demultiplexing and linkers trimming using FASTX-toolKit (0.0.13). Clean CAGE tags were aligned to the mouse reference genome (mm10) using Bowtie2 with the -- very-sensitive preset (set as -D 20 -R 3 -N 0 -L 20 -i S,1,0.50), and the resulting SAM files were converted to sorted BAM files using SAMTools (1.9). BAM files of Light and Heavy polysome fractions were merged to form one BAM file representing all translated fractions, marked as Polysomes. BAM files were then loaded into the CAGEr package (v1.32 in the Bioconductor environment, v3.12) and uniquely-mapped tags were used for calling CAGE-defined TSSs (CTSSs) setting parameters as follows: sequencingQualityThreshold = 20 and mappingQualityThreshold = 20. As part of CTSS mapping, CAGEr performed corrections for the CAGE-specific G-bias caused by template-free guanine incorporation upstream to the true transcription start site by the reverse transcriptase used for cDNA synthesis ^3^. Metagene analysis for profiling CTSS coverage was performed using the ’metagene’ package in the Bioconductor environment. Eµ-Tcl1 and WT CTSS locations were compared to the reference dataset for transcription tart sites (refTSS) of the mouse (v3.1) published by RIKEN institute (46).

### CAGE tag clustering, quantification and analysis

Most of the CTSS analysis was performed using CAGEfightR package (1.10.0) 4, which uses several R-packages from the Bioconductor project, v3.12. BIGWIG files that were exported by the CAGEr were loaded into CAGEfightR and closely-spaced CAGE tags were clustered into unidirectional and bidirectional tag clusters (TCs) using the slice-reduce approach to define the global TSSs and enhancer candidates, respectively. TCs with less than one Tag-Per-Million (TPM) in the two Eµ-Tcl1 or WT CAGE libraries were removed in further analysis. Active enhancers predicated as balanced (threshold scored as 0.95) bidirectional transcription of capped enhancer RNAs (eRNAs). TSS-enhancer physical interactions predicted by distance (50kbp) and correlation (Pearson > 0, *P*<0.05) as previously shown by Andersson et al., 2014. TxIDs, with their structure models (e.g., promoter/UTR/CDS/exon/intron/antisense) and GeneID were assigned to each TC using the two genome-wide annotation packages, ’TxDb.Mmusculus.UCSC.mm10.knownGene’ (47) and ‘org.Mm.eg.db’ (48), of the Bioconductor project, respectively. TC widths measured between 10-90% of the interquartile range (IQR) of pooled CAGE tags and resulted in a bimodal distribution of sharp (1-10bp) and broad (11-100 bp) TCs to be analyzed separately further in the analysis. Cluster-level and Gene-level differential expression analyses were performed by Deseq2 and limma packages of the Bioconductor project. Differential TSS Usage (DTU) analysis was performed using the edgeR diffSpliceDGE method in a subset of genes with multi TSSs corresponding to more than 10% of total gene expression.

### Motif-based sequence analysis for TF binding sites

In order to know what transcription factors might be involved in the regulation of alternative TSSs, we downloaded DNA-binding motifs as position frequency matrices (PFMs) from the core collection of JASPAR database (49) and matched it against promoter regions defined as -1000 bp and +100 to the peaks of CAGE tag clusters of TSSs and enhancers. We used TFBSTools (50) and motifmatcher (51). motifmatchr: Fast Motif Matching in R) packages of the Bioconductor project, to obtain and find matches of motifs PFMs in promoter sequences, respectively. Using Fisher’s Exact test, we have been able to see if a motif significantly co-occurs in upregulated alternative TSS and enhancers.

### Eµ-Tcl1 adoptive transfer model

Generation of this mouse model was performed as previously described (52). Briefly, Eµ-Tcl1 mice approximately 12 months of age, with a malignant cell population higher than 60% in the peripheral blood (PB) were sacrificed. Their spleens were excised, and 4 × 10^7 cells resuspended in PBS-/- were injected into the tail vein of 6–8-week-old WT recipient mice. Progression of the disease was followed in the PB by using flow cytometry for the IgM/CD5 population. Mice with >30% IgM+/CD5+ cells were considered to be diseased and were used for further analysis.

### Inhibitor assay

The survival of CLL B cells and healthy B cells were tested under the inhibitory effect of drugs targeting the activity of HDAC type II (TMP269, Cayman cat. 17738), HDAC type III (Thiomyristoyl, Cayman cat. 19398), KDM4A (NSC636819, Sigma-Aldrich, cat. 531996), DNMT3A (Decitabine, MCE cat. HY-A0004) and EIF4E (Tomivosertib, Cayman cat. 21957). B cells isolated from Eu-Tcl1 adoptive transfer and WT mice as described above. Triplicates of 2 million cells (per well) distributed in a 24-well tissue-culture plate treated with a serial four incremented concentrations of the above inhibitors (in DMSO). Following 48h, Cells were washed with cold PBS, and resuspended in Annexin V Binding Buffer (BioLegend) at a concentration of 0.25-1.0 x 10^7 cells/ml. FITC Annexin V (BDPharmingen cat. 556419) was added to the samples with anti CD19 and anti CD5 for 20 minutes in the dark on ice. Cells were washed twice with Annexin V Binding Buffer (centrifuge 1400rpm for 5 minutes, discard the supernatant), then resuspended in 300µL of Annexin V Binding Buffer with 7- AAD (BDparmingen cat. 88981E) Viability Staining Solution. Cells immediately were analyzed by Flow Cytometry.

### Data availability

The CAGE-seq and Polysome-CAGE data generated in this study were deposited in NCBI’s Gene Expression Omnibus and will be made public upon acceptance of the manuscript.

## Author Contributions

R.D. conceived the study; R.D., T.H.S, A.O. and I.S designed the study; A.O., T.H.S., S.H.B. and K.D. carried out the experiments; A.O. and R.D. analyzed the data and wrote the paper.

## Supplementary figure legends

**Figure 1- figure supplement 1.**
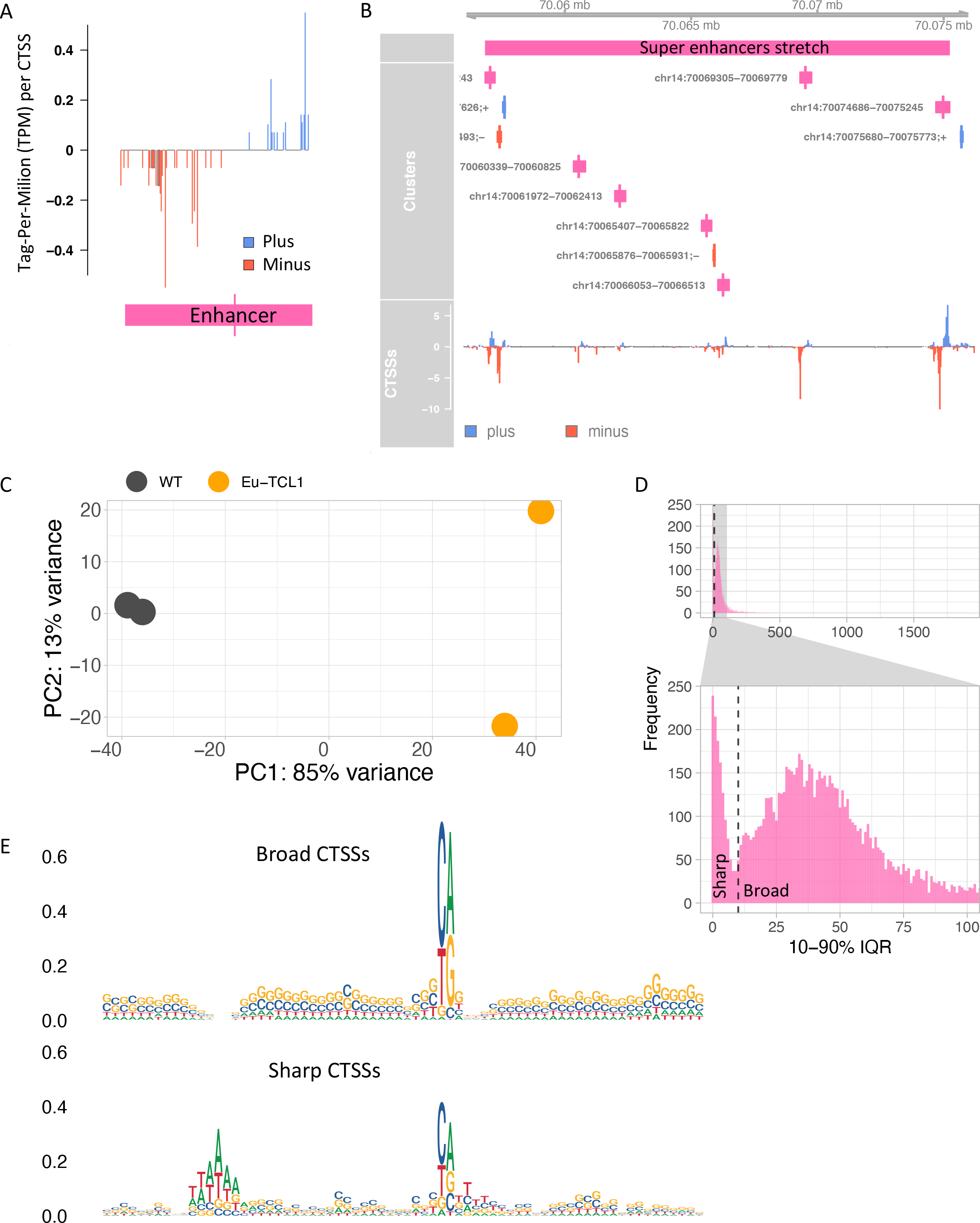
A. Genome browser example of an enhancer candidate showing balanced bidirectional transcription of enhancer RNAs (eRNAs). Tag-Per-Million per values plotted in blue for plus strand and in red for minus strand. **B.** An example of a super enhancer. Closely spaced enhancers positioned within 18,405 bps range. **C.** “Blind” version of the variance-stabilizing transformation of the two replicates of Eu-Tcl1 and WT CAGE samples. **D.** Bimodal distribution of the widths of tag clusters of highly expressed TSSs. Most CAGE tag clusters (TC) are distributed either below (Sharp) or above (Broad) 10 bp distance holding 10-90% of pooled CAGE tags. **E.**Sequence logos of core promoter regions (-40 and +30 bp) of Sharp and Broad classes of TCs.

**Figure 2- figure supplement 1.**
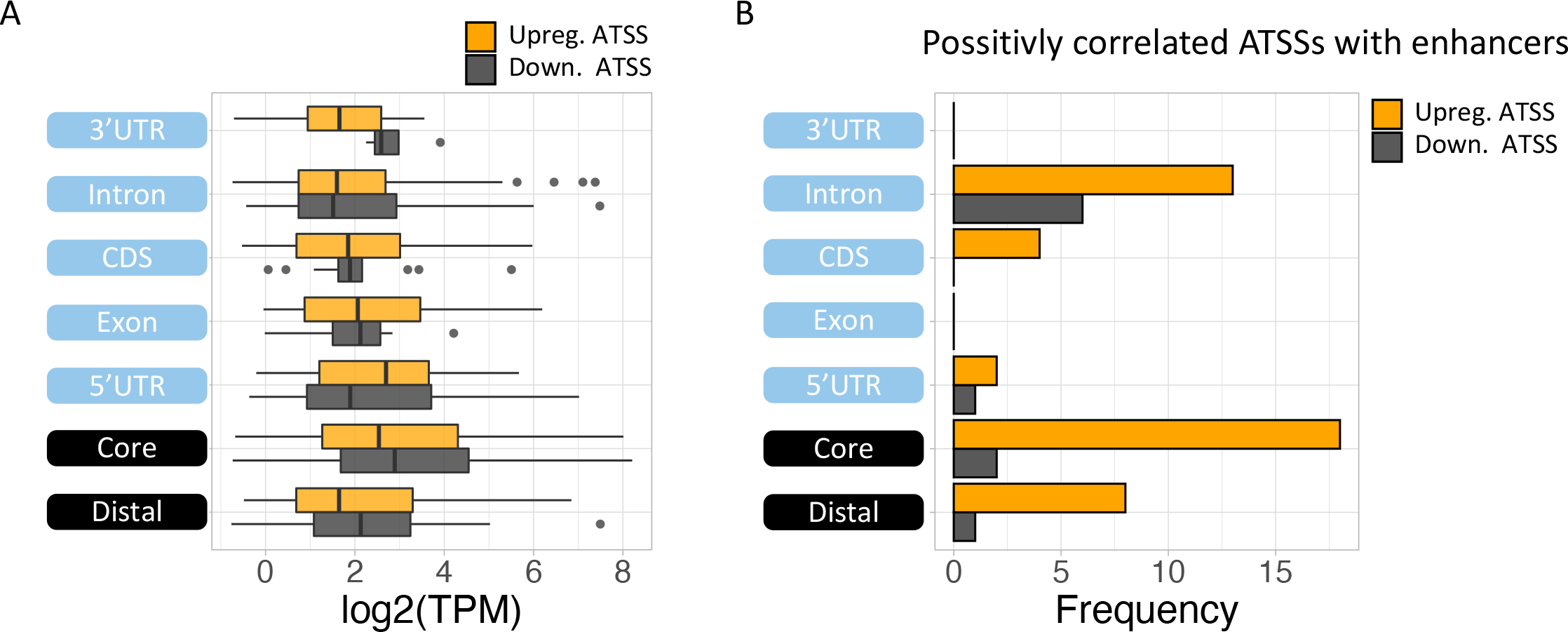
A. Boxplot of expression level (log2(TPM)) of alternative TSS (ATSS) upregulated (orange) or downregulated (grey) in Eu-Tcl1 by gene-structure location. **B.** The frequency of initiating 5’ UTR single base of Eu-Tcl1 (orange) and WT (grey) TSSs. **C.** Frequency of initiating 5’UTR trinucleotides in overall TSSs of Eu-Tcl1 (orange bars) and WT (grey bars), scaled in the left Y axis and in upregulated ATSS (orange line) and downregulated ATSS (grey line), scaled in the right Y axis. **D.** The frequency of positively correlated ATSSs with active enhancers by gene-structure location.

**Figure 5- figure supplement 1.**
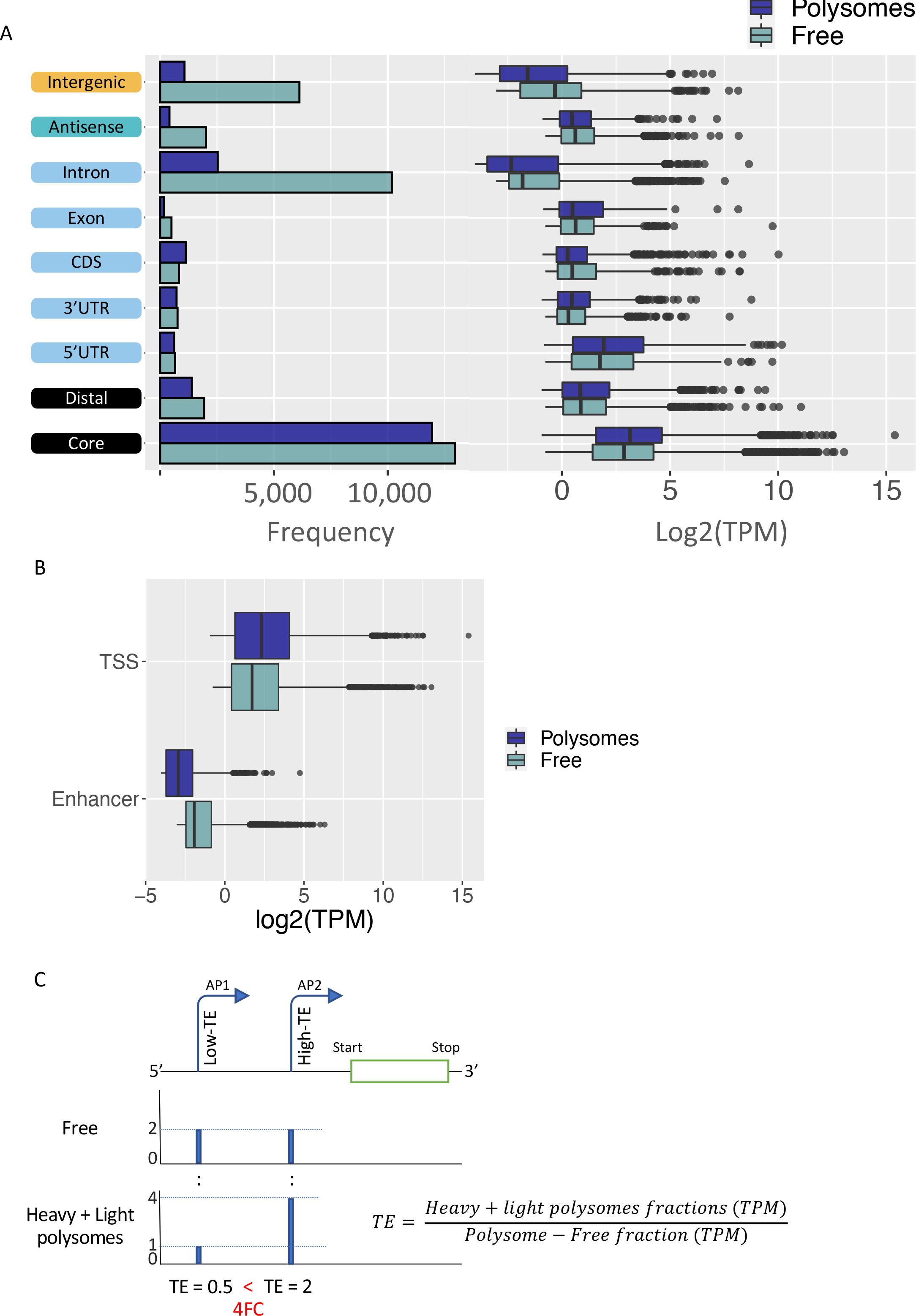
A. Frequency and boxplot of expression level (log2(TPM)) of of CAGE tag clusters in polysome-free (light blue) and Heavy polysomes (dark blue) by gene-structure locations. **B.** Boxplot of expression level (log2(TPM)) of tag clusters classified as TSS or enhancer in polysome-free (light blue) and Heavy polysomes (dark blue). **C.** Example of differentially translated isoforms originated by alternative promoters (P1 and P2), differing in 5’UTR lengths. TSS coverage tracks of polysome-free (Free) and Heavy + Light polysomes are used for calculating TE values of the alternative isoforms. When the TE of the one isoform (shorter 5’UTR, AP2) is 2FC higher than the longer isoform (AP1), the first assigned as Low-TE isoform and the latter as High-TE isoform.

**Figure 6- figure supplement 1.**
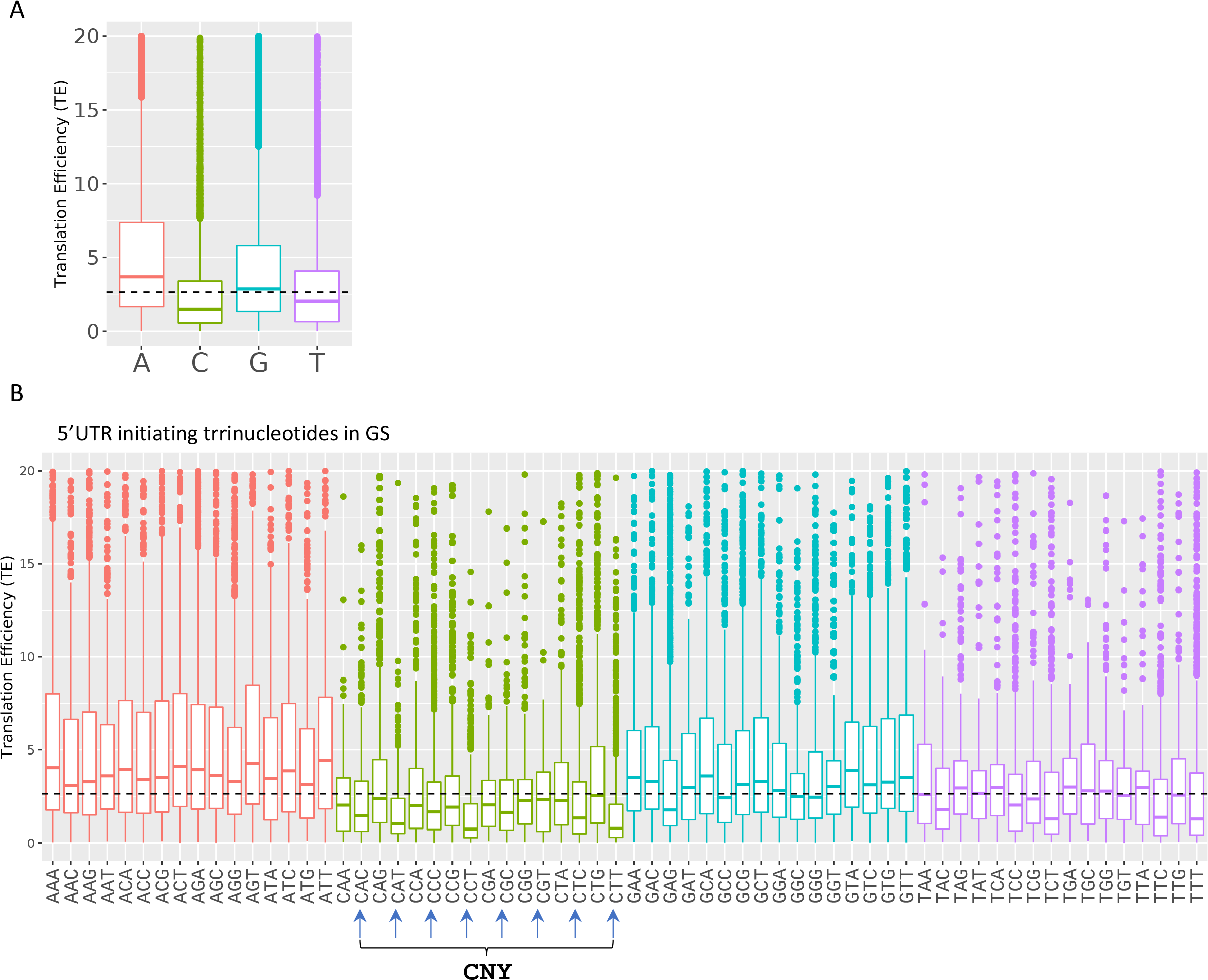
A-B. Boxplots of transcription efficiency (TE) of isoforms differ in their initiating TSS single nt (A) and first three nucleotides (B) are distributed and colored by initiating Adenosine, Cytosine, Guanin and Tyrosine. (C) Horizonal line represents overall median of TE values. All the data presented in this figure are the mean of the two independent replicates. The bottom and the top whiskers represent 5% and 95% of the distribution, respectively.

### Supplementary tables

**Figure 2 supplementary table 1.**
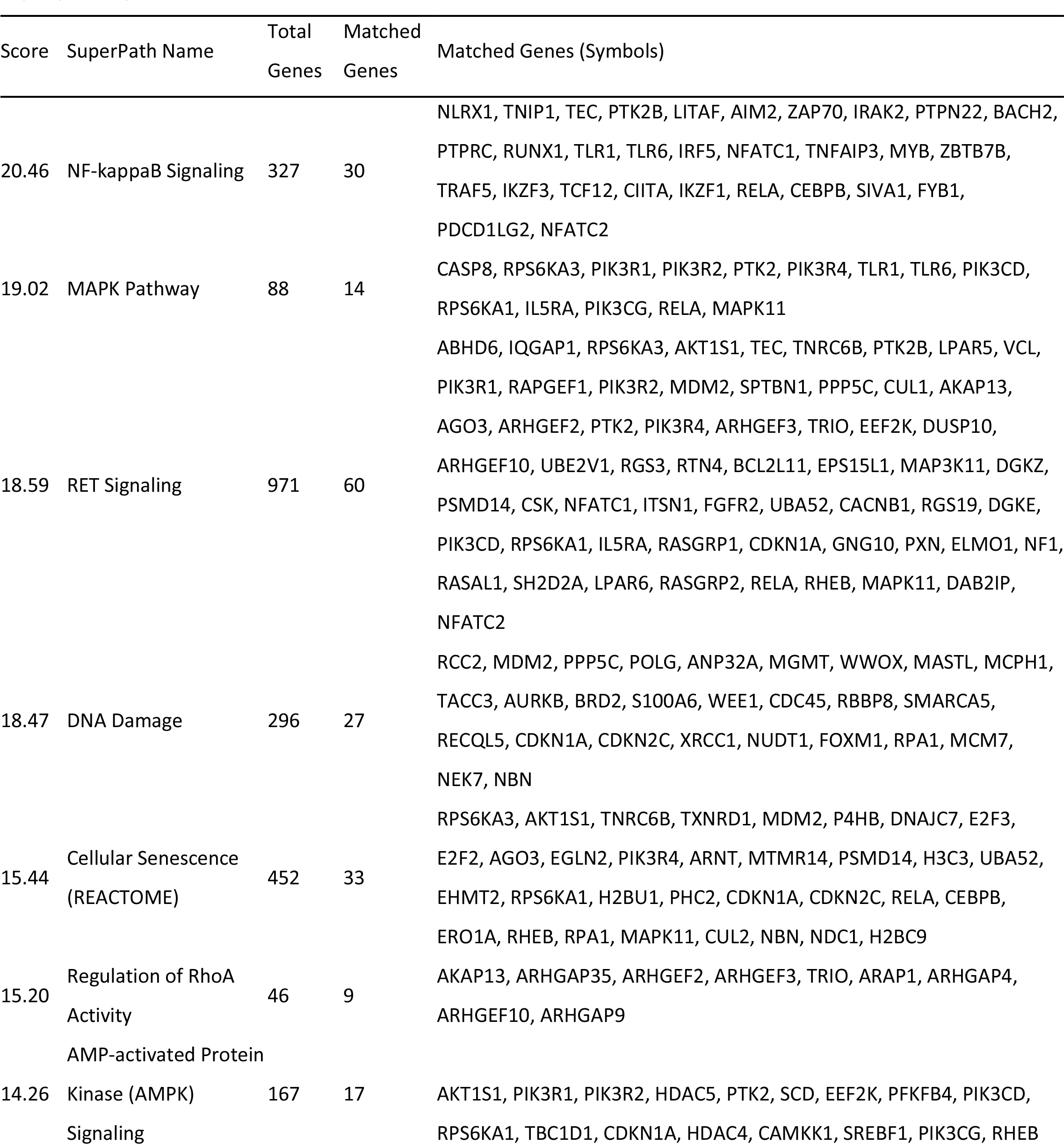
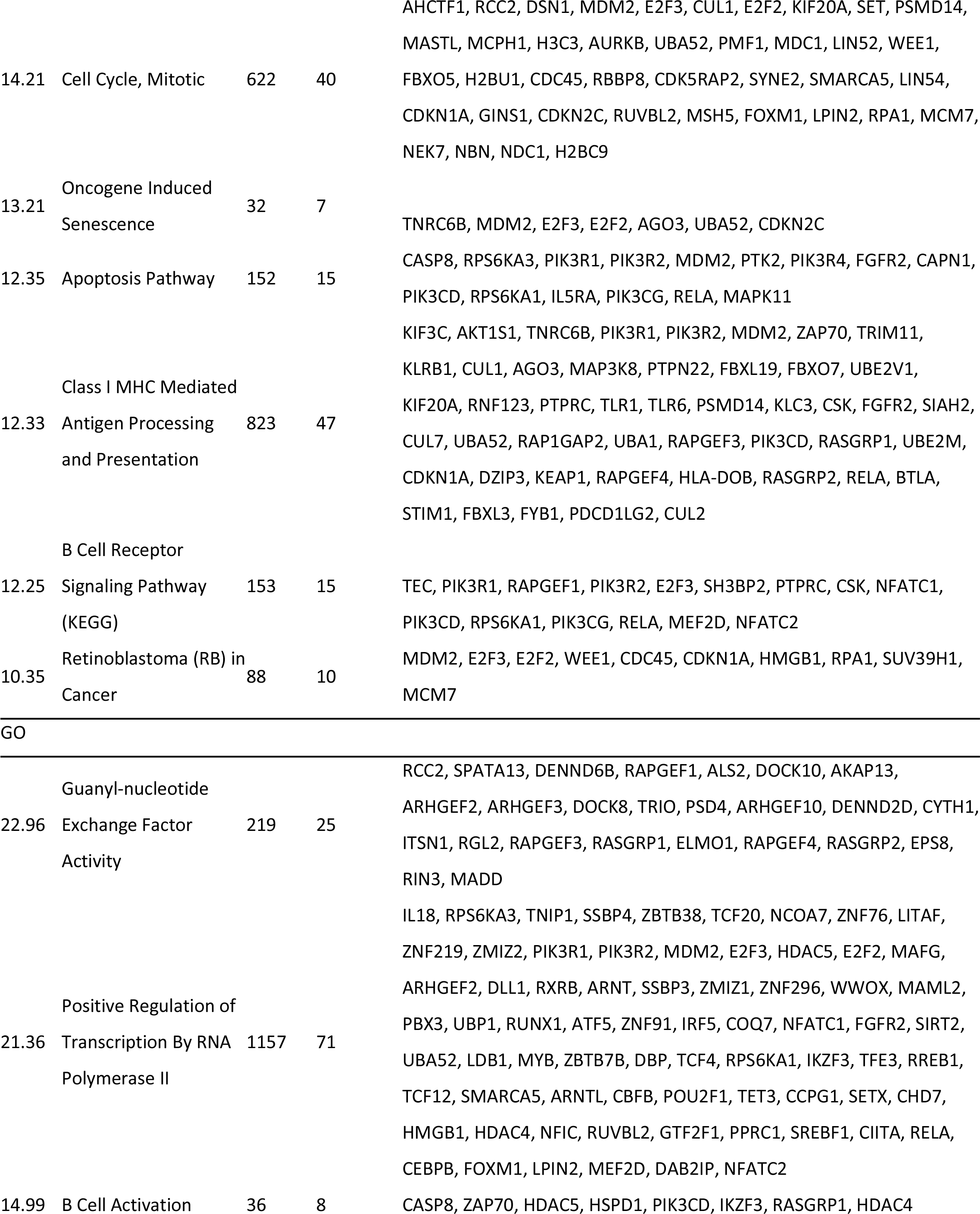
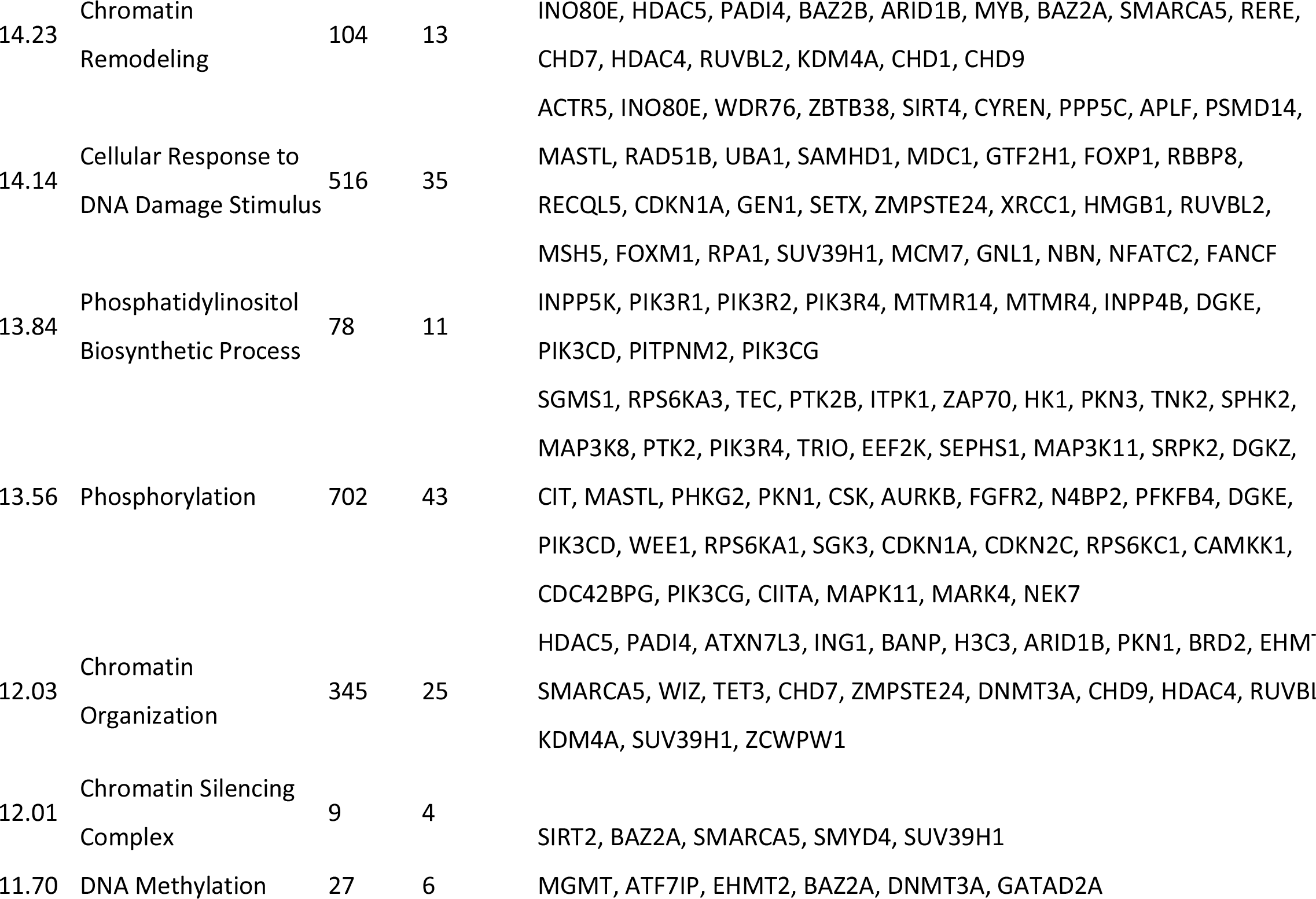
GO pathways affected by differential TSS usage. Super-pathways

**Figure 2 supplementary table 2.** MGI phenotypes affected by differential TSS usage.

| Score | Name | Total<br>Genes | Matched<br>Genes | Symbols |
| --- | --- | --- | --- | --- |
| 31.00 | Increased B Cell Number | 92 | 19 | TNIP1, APOE, ZAP70, SPHK2, PTPN22, DLL1, RGS3, BCL2L11, PTPRC, DGKZ, MCPH1, TNFAIP3, TNFRSF13B, RASGRP1, CDKN1A, CDKN2C, IKZF1, SEMA4D, NFATC2 |
| 20.68 | Thymus Hypoplasia | 118 | 17 | WIPF1, AFF1, ZFP36, MDM2, ZAP70, NDST1, PTPRC, RUNX1, TCF12, CBFB, CHD7, PNP, ZMPSTE24, PIK3CG, IKZF1, NBN, GFI1 |
| 19.13 | Enlarged Spleen | 254 | 25 | TNIP1, ZFP36, MDM2, PPP5C, E2F2, PTPN22, ING1, WWOX, BCL2L11, PTPRC, RUNX1, PSTPIP2, UNC13D, TNFAIP3, MYB, TNFRSF13B, IKZF3, CBFA2T3, RASGRP1, CDKN1A, CDKN2C, IKZF1, CEBPB, STIM1, NFATC2 |
| 17.97 | "decreased CD4-positive,<br>Alpha Beta T Cell<br>Number" | 136 | 17 | ZAP70, DLL1, DOCK8, FBXO7, PTPRC, RUNX1, DGKZ, SIRT2, ZBTB7B, IL5RA, RASGRP1, PNP, RUVBL2, PIK3CG, CIITA, SEMA4D, NBN |
| 16.05 | Abnormal Cytokine<br>Secretion | 151 | 17 | LITAF, AIM2, MAP3K8, SEMA4A, DUSP10, PTPRC, TLR1, TLR6, PSTPIP2, UNC13D, TNFAIP3, ZBTB32, CIITA, NLRC4, STIM1, PDCD1LG2, GFI1 |

**Figure 4 supplementary table 1.**
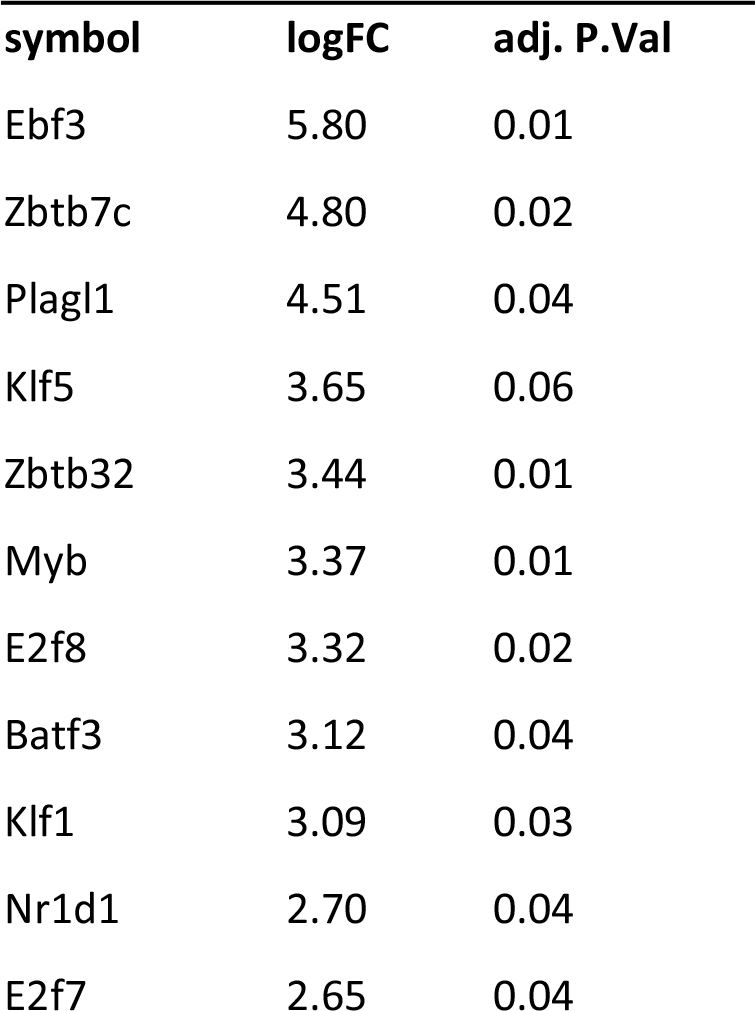
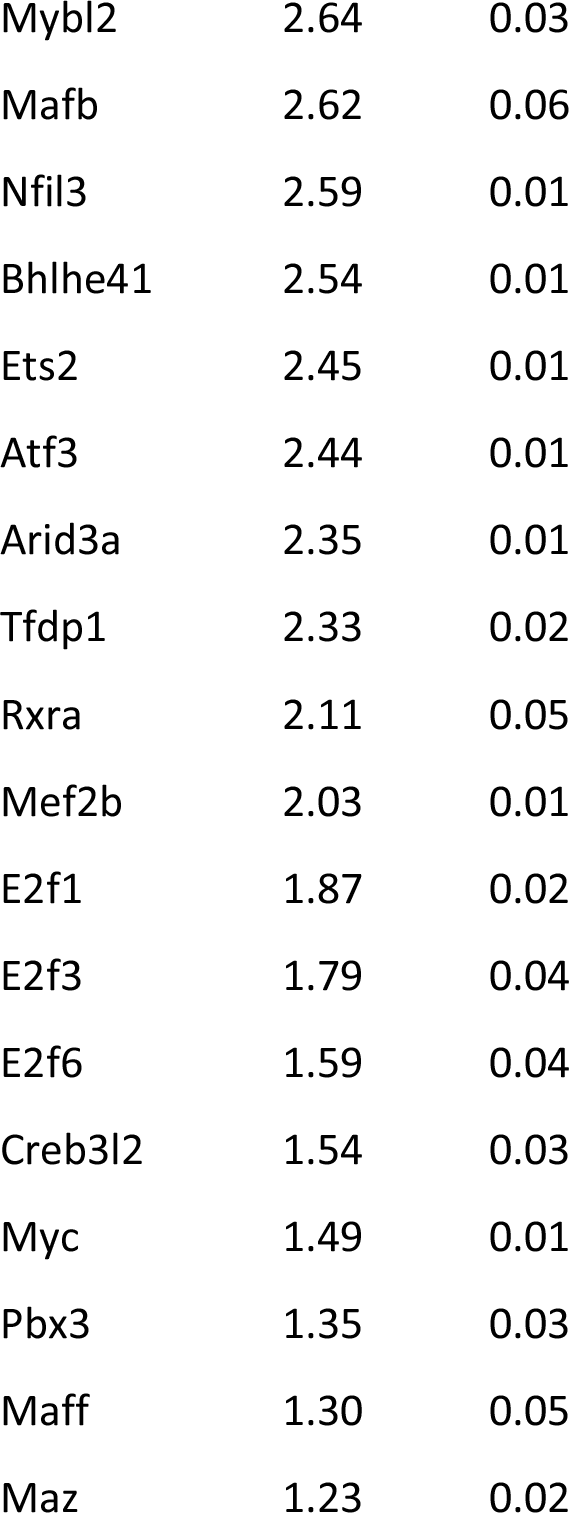
Transcription factors upregulated in Eu-Tcl1

